# Spot Dynamics in a Reaction-Diffusion Model of Plant Root Hair Initiation

**DOI:** 10.1101/114876

**Authors:** Daniele Avitabile, Victor F. Breña-Medina, Michael J. Ward

## Abstract

We study pattern formation in a 2-D reaction-diffusion (RD) sub-cellular model characterizing the effect of a spatial gradient of a plant hormone distribution on a family of G-proteins associated with root-hair (RH) initiation in the plant cell *Arabidopsis thaliana*. The activation of these G-proteins, known as the Rho of Plants (ROPs), by the plant hormone auxin, is known to promote certain protuberances on root hair cells, which are crucial for both anchorage and the uptake of nutrients from the soil. Our mathematical model for the activation of ROPs by the auxin gradient is an extension of the model of Payne and Grierson [PLoS ONE, **12**(4), (2009)], and consists of a two-component Schnakenberg-type RD system with spatially heterogeneous coefficients on a 2-D domain. The nonlinear kinetics in this RD system model the nonlinear interactions between the active and inactive forms of ROPs. By using a singular perturbation analysis to study 2-D localized spatial patterns of active ROPs, it is shown that the spatial variations in the nonlinear reaction kinetics, due to the auxin gradient, lead to a slow spatial alignment of the localized regions of active ROPs along the longitudinal midline of the plant cell. Numerical bifurcation analysis, together with time-dependent numerical simulations of the RD system are used to illustrate both 2-D localized patterns in the model, and the spatial alignment of localized structures.

## 1. Introduction

We examine the effect of a spatially-dependent plant hormone distribution on a family of proteins associated with root hair (RH) initiation in a specific plant cell. This process is modeled by a generalized Schnakenberg reaction-diffusion (RD) system on a 2-D domain with both source and loss terms, and with a spatial gradient modeling the spatially inhomogeneous distribution of the plant hormone auxin. This system is an extension of a model proposed by Payne and Grierson in [33], and analyzed in a 1-D context in the companion articles [4, 6]. The new goal of this paper, in comparison with [4, 6], is to analyze 2-D localized spot patterns in the RD system (1.2), and how these 2-D patterns are influenced by the spatially inhomogeneous auxin distribution.

We now give a brief description of the biology underlying the RD model. In this model, an on-and-off switching process of a small G-protein subfamily, called the Rho of Plants (ROPs), is assumed to occur in a RH cell of the plant *Arabidopsis thaliana*. ROPs are known to be involved in RH cell morphogenesis at several distinct stages (see [13, 23] for details). Such a biochemical process is believed to be catalyzed by a plant hormone called auxin (cf. [33]). Typically, auxin-transport models are formulated to study polarization events between cells (cf. [14]). However, little is known about specific details of auxin flow within a cell. In [4, 6, 33] a simple transport process is assumed to govern the auxin flux through a RH cell, which postulates that auxin diffuses much faster than ROPs in the cell, owing partially to the in-and out-pump mechanism that injects auxin into RH cells from both RH and non-RH cells (cf. [16, 18]). Recently, in [27], a model based on the one proposed in [33] has been used to describe patch location of ROP GTPases activation along a 2-D root epidermal cell plasma membrane. This model is formulated as a two-stage process, one for ROP dynamics and another for auxin dynamics. The latter process is assumed to be described by a constant auxin production at the source together with passive diffusion and a constant auxin bulk degradation away from the source. In [27] the auxin gradient is included in the ROP finite-element numerical simulation after a steady-state is attained. Since we are primarily interested in the biochemical interactions that promote RHs and key ingredients that geometrical features have on the RHs initiation dynamics, rather than providing a detailed model of auxin transport in the cell, we shall hypothesize specific, but qualitatively reasonable in the sense described below, time-independent spatially varying expressions for the auxin distribution in a 2-D azimuthal projection of a 3-D RH cell wall. In this way, we capture crucial ingredients of the spatially-dependent auxin distribution in the cell. Qualitatively, since a RH cell is an epidermal cell, influx and efflux biochemical gates are distributed along the cell membrane, and the auxin flux is known to be primarily directed from the tip of the cell towards the elongation zone (cf. [16, 21]), leading to a longitudinally spatially decreasing auxin distribution. Our specific form for the auxin distribution will incorporate such a longitudinal spatial dependence. However, as an extension to the auxin model developed in [4, 6], here we also can take into account non-RH lateral contributions of auxin flux in RH cells by considering an auxin distribution with a transverse spatial dependence. This assumption arises as non-RH cells are believed to promote auxin transversal flux into RH cells. However, although auxin is believed to follow pathways either through the cytoplasm or through cell walls of adjacent cells, we are primarily interested in the former pathway that is directed within the cell, and which induces the biochemical switching process (cf. [31]). In this new 2-D analysis of the RD model, we will allow for an auxin distribution that has an arbitrary spatial dependence. However, in illustrating our 2-D theory, and motivated by the biological framework discussed above, we will focus on two specific forms for the auxin distribution, as shown in Figure 1: a 1-D form that has the spatially decreasing longitudinal dependence of [4, 6], and a modification of this 1-D form that allows for a transverse dependence. We will show that these specific auxin gradients induce an alignment of localized regions of elevated ROP concentration, referred to here as spots. The RD system under consideration exhibits spot-pinning phenomena, in the sense that the position of spots in the stationary patterns is determined by the spatial gradient of auxin. As we shall see below, spots slowly drift until they become aligned with the longitudinal axis of the cell. Their steady-state locations along this cell-midline are at locations determined by the auxin gradient, which effectively “pins” the spots. A similar pinning phenomenon is found for spatially localized structures in other systems: an example is the vortex-pinning phenomenon in the Ginzburg–Landau model of superconductivity in an heterogeneous medium, see [1].

**Figure 1:**
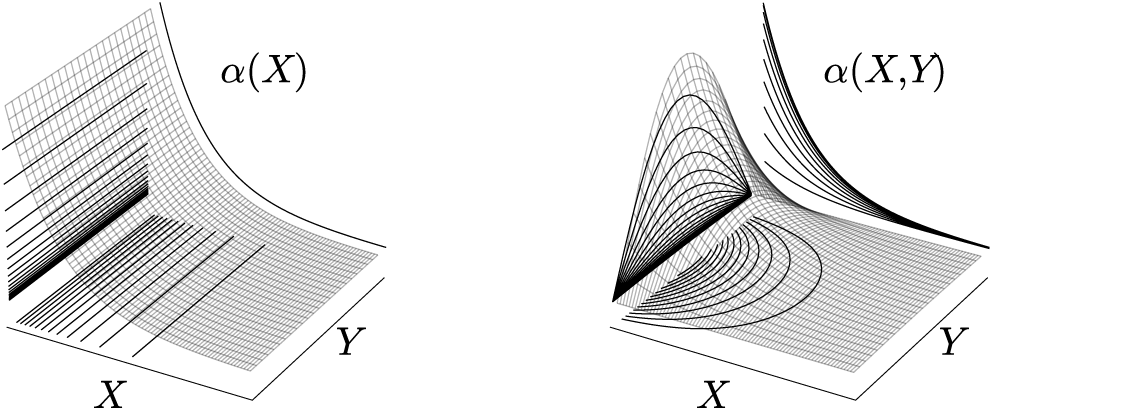
*Sketches of the spatially-dependent auxin gradient forms: constant in the transverse direction (left-hand panel), and decreasing on either side of the middle of the cell (right-hand panel).*

In Figure 2, a sketch of an idealized 3-D RH cell is shown. In this figure, heavily dashed gray lines represent the RH cell membrane. The auxin flux is represented by black arrows, which schematically illustrates a longitudinal decreasing auxin pathway through adjacent cells. As ROP activation is assumed to occur near the cell wall and transversal curvature of the domain is not included in our model, we consider a projection onto a 2-D rectangular domain of this cell, as is also shown in Figure 2.

**Figure 2:**
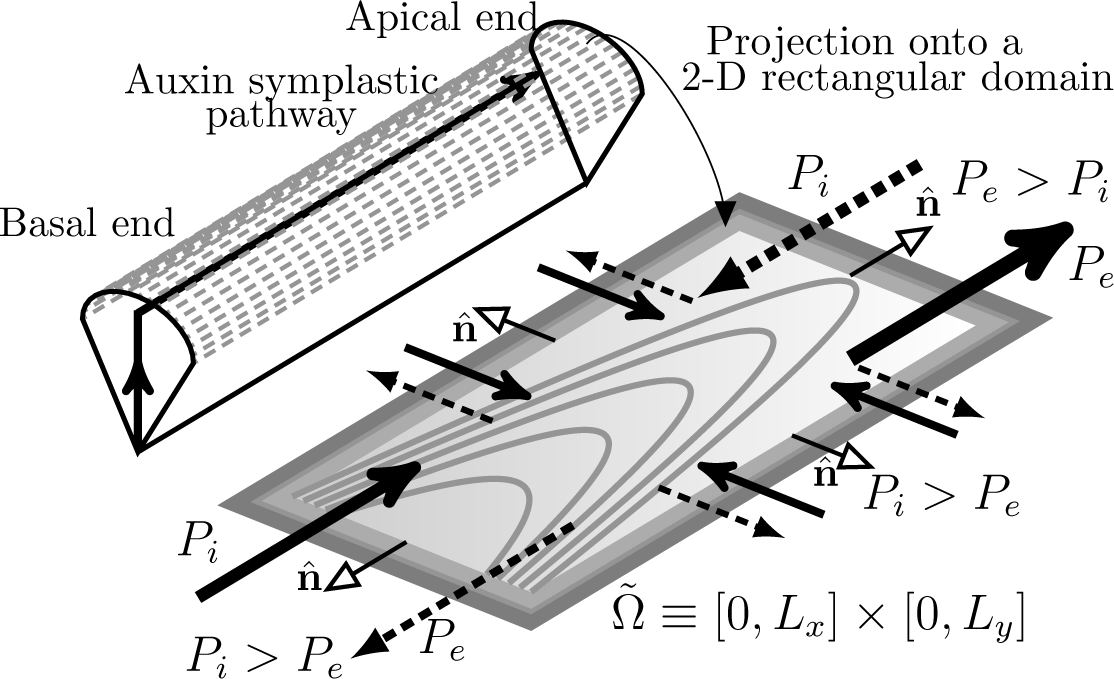
*Sketch of an idealized 3-D RH cell with longitudinal and transversal spatially-dependent auxin flow. The auxin gradient is shown as a consequence of in- and out-pump mechanisms. Influx P*_*i*_ *and efflux P_e_ permeabilities are respectively depicted by the direction of the arrow; the auxin symplastic pathway, indicated by black solid arrows in the 3-D RH cell, is the auxin pathway between cells. Here, the cell membrane is depicted by heavily dashed gray lines. The gradient is coloured as a light gray shade, and auxin gradient level curves are plotted in gray. Cell wall and cell membrane are coloured in dark and light gray respectively. Modified figure reproduced from [6].*

In [6] we considered the 1-D version of the RD model given in (1.2), where the RH cell is assumed to be slim and flat, and where the auxin distribution depends only on the longitudinal spatial direction. There it was shown that 1-D localized steady-state patterns, representing active-ROP proteins, slowly drift towards the interior of the domain and asymmetrically get pinned to specific spatial locations by the auxin gradient. This pinning effect, induced by the spatial distribution of auxin, has been experimentally observed in, for instance, [13, 15]. Biologically, although multiple swelling formation in transgenic RH cells may occur, not all of these will undergo the transition to tip growth (cf. [22]). In [6] a linear stability analysis showed that multiple-active-ROP 1-D spikes may be linearly stable on 𝒪(1) time-scales to 1-D perturbations, with the stability properties depending on an inversely proportional relationship between length and auxin catalytic strength. This dynamic phenomenon is a consequence of an auxin catalytic process sustaining cell wall softening as the RH cell grows. For further discussion of this correlation between growth and biochemical catalysis see [6].

In [4] the linear stability of 1-D stripe patterns of elevated ROP concentration to 2-D spatially transverse perturbations was analyzed. There it was shown that single interior stripes and multi-stripe patterns are unstable under small transverse perturbations. The loss of stability of the stripe pattern was shown to lead to more intricate patterns consisting of spots, or a boundary stripe together with multiple spots (see Figure 9 of [4]). Moreover, it was shown that a single boundary stripe can also lose stability and lead to spot formation in the small diffusivity ratio limit.

**Figure 3:**
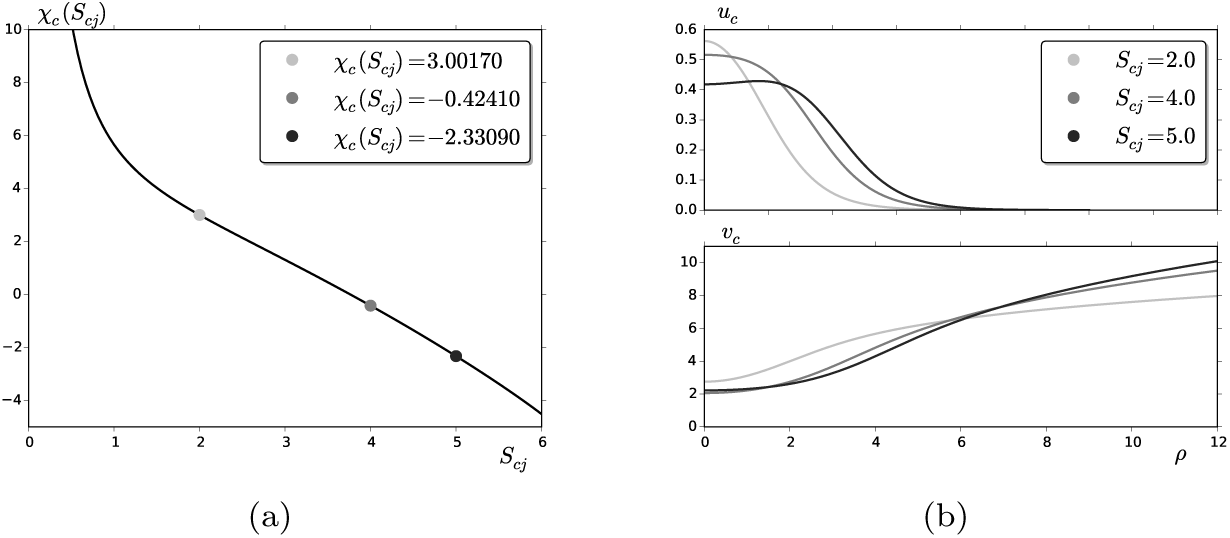
*(a) The constant χ*_*c*_ (*S*_*cj*_) *versus the source parameter S*_*cj*_, *computed numerically from* (2.14). *(b) Radially symmetric solution u_c_ (upper panel) and v_c_ (bottom panel), respectively, at the values of S*_*cj*_ *shown in the legend. Parameter Set A from Table 1, where τ/β* = 3.

**Figure 4:**
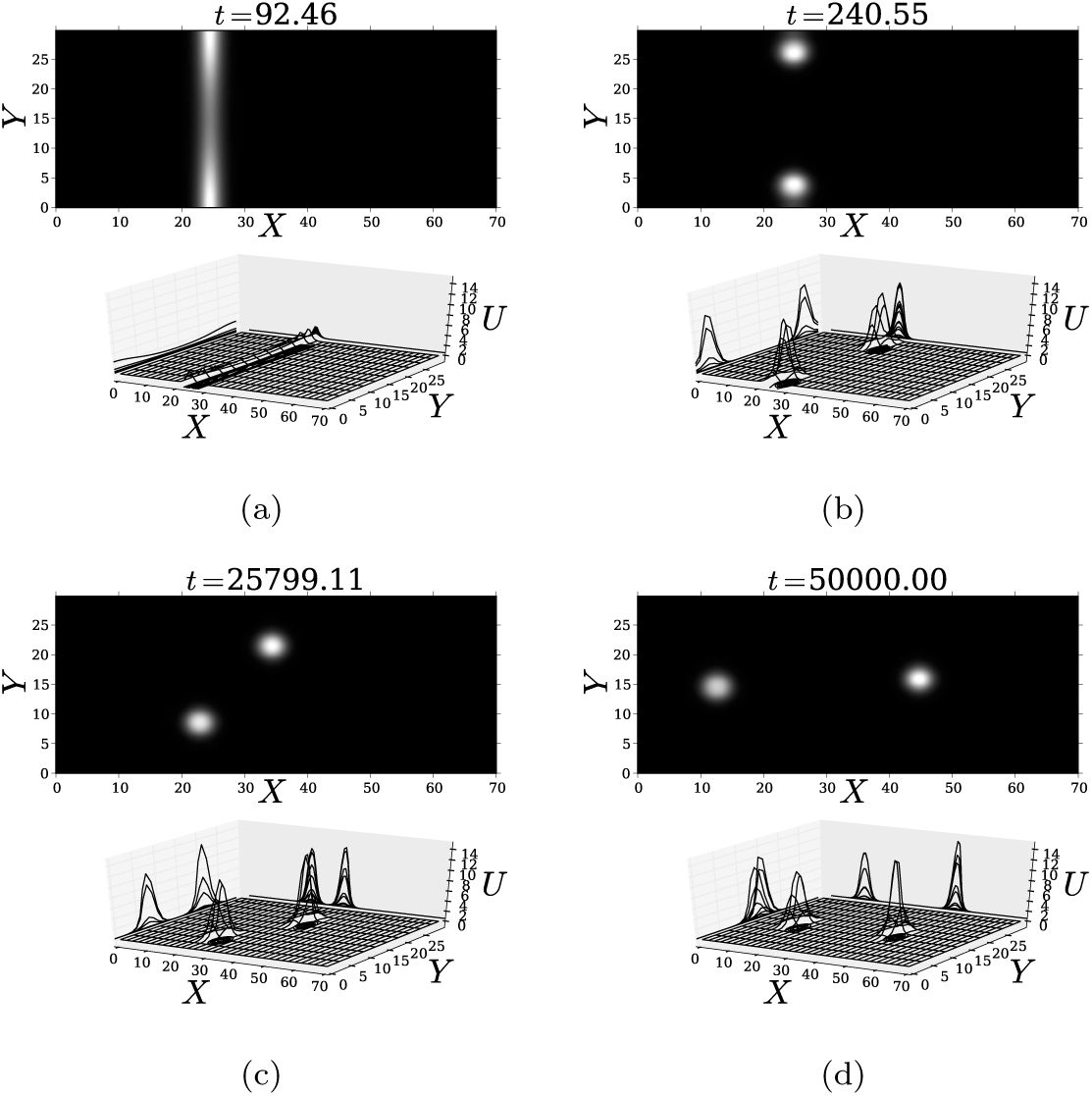
*Breakup instability of an interior localized homoclinic stripe for U, and the resulting slow spot dynamics. (a) The localized stripe initially breaks into two spots; (b) once formed, the spots migrate from the boundary towards each other in the vertical direction, and then (c) rotate until they get aligned with the longitudinal direction. (d) Finally, they get pinned far from each other along the y-midline by the auxin gradient. Parameter Set A from Table 1 was used with k*_20_ = 0.5. *For these values the initial steady-state stripe is centered at X*_0_ = 24.5. *Figure from [4].*

**Figure 5:**
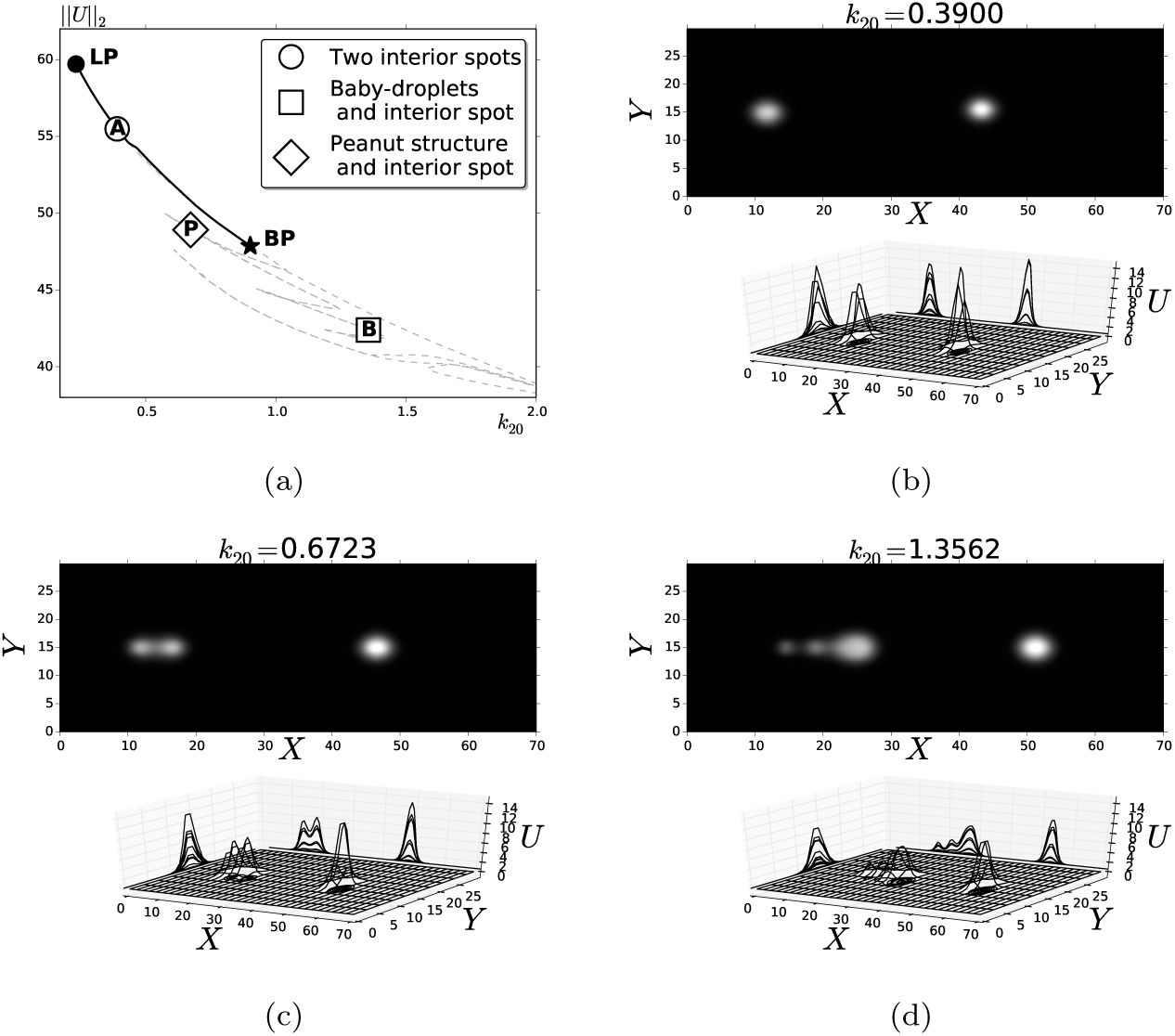
*(a) Bifurcation diagram as k*_20_ *is varied showing a stable branch, labelled A, of (b) two* 20 *spots, and unstable branches (light gray-dashed curves). Filled circle at 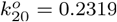 is a fold bifurcation (LP) and the filled star at 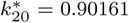is a pitchfork bifurcation (BP). (c) A peanut structure and one spot steady-state, from label P in (a). (d) A two-spot with baby droplets steady-state solution, from label B in (a). Parameter Set A was used, as given in Table 1. The k*_20_ *values are shown on top of upper panels in (b)-(d).*

**Figure 6:**
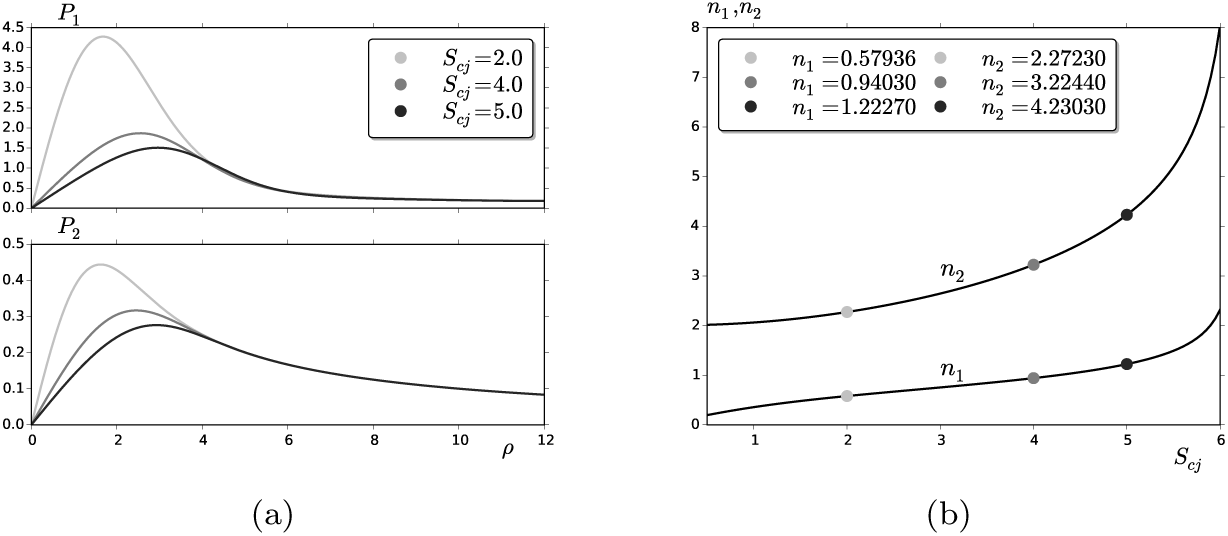
*(a) Numerical solution of the adjoint solution satisfying* (3.31) *for* **P** = (*P*_1_, *P*_2_)^*T*^ *for values of S*_*cj*_ *as shown in the legend; P*_1_ *(top panel) and P*_2_ *(bottom panel). (b) Numerical results for the solvability condition integrals n*_1_ *and n*_2_, *defined in* (3.34b), *as S*_*cj*_ *increases. The filled circles correspond to S*_*cj*_ *values in (a). We use Parameter Set A from Table 1, for which τ/β* = 3.

**Figure 7:**
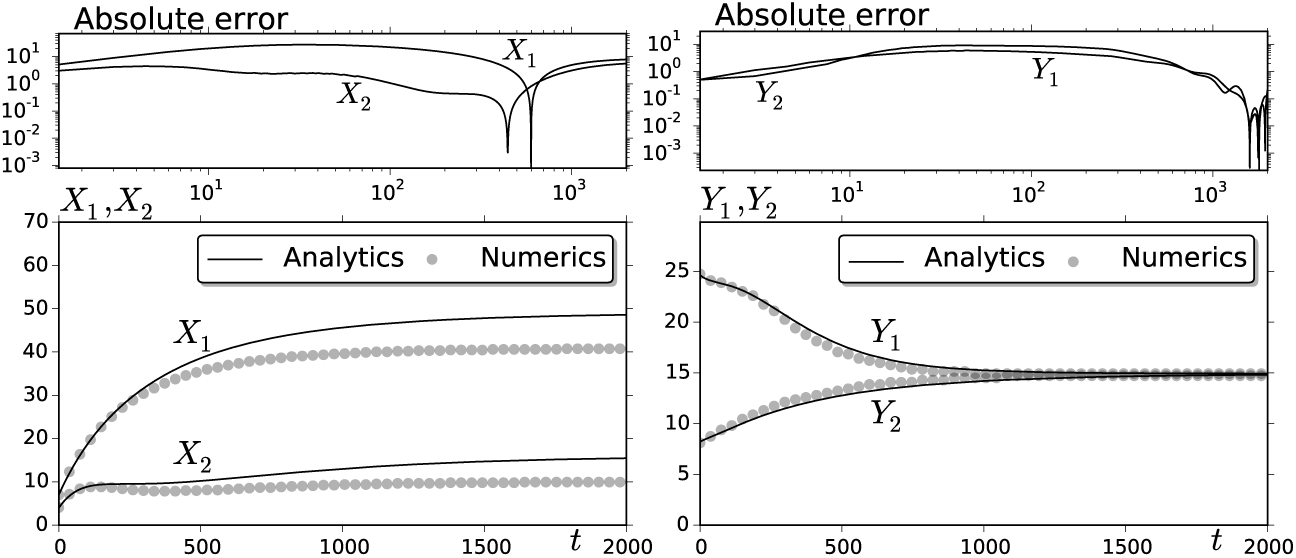
*The time-dependent location of two spots for an auxin gradient of the type (i) in* (1.1) *as obtained from the DAE system (solid curves) in Proposition 2 with k*_20_ = 0.3 *and with parameters of Parameter Set A of Table 1. The initial locations for the two spots are X*_1_(0) = 7.0, *Y*_1_(0) = 8.21 *and X*_2_(0) = 3.97, *Y*_2_(0) = 24.63. *The full numerical results computed from* (1.2) *are the filled circles; absolute error in the top panels. Observe that the DAE system predicts that the two spots become aligned along the longitudinal midline of the cell. The x-coordinate (left-hand panels) and the y-coordinate (right-hand panels) of the two spots.*

**Figure 8:**
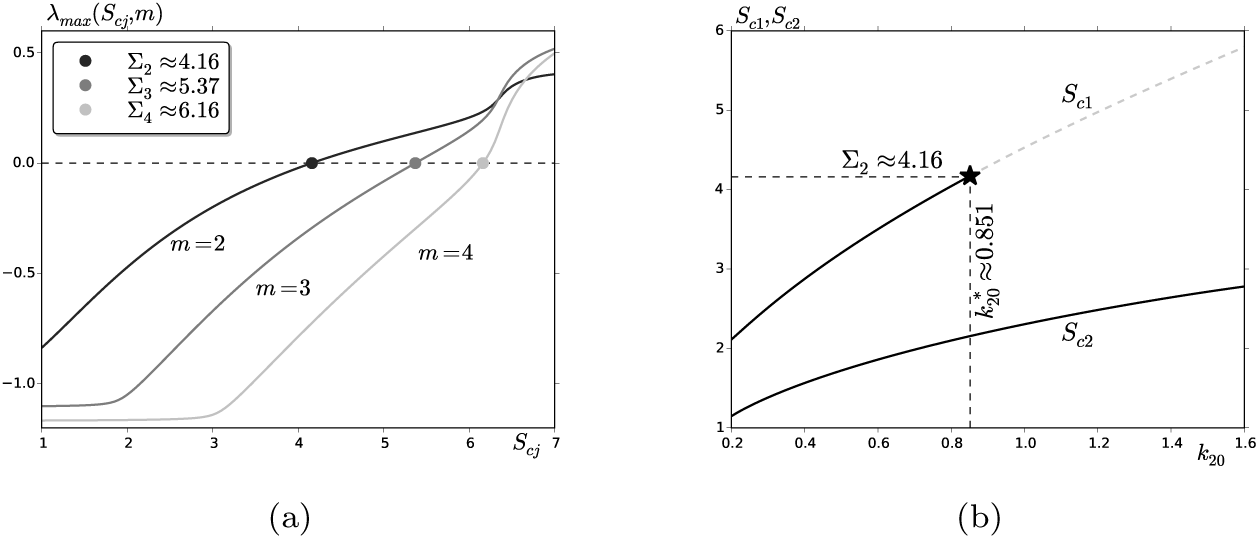
*(a) The largest real-valued eigenvalue λ*_*max*_ *of* (4.37) *versus S*_*cj*_ *for different angular modes m. The peanut-splitting thresholds Σ_m_ for m = 2,3,4 are indicated by filled circles. (b) Source parameters for a two-spot steady-state solution of the DAE system* (3.34) *as k*_20_ *is varied. Stable profiles are the solid curves, and the spot closest to the left boundary at x* = 0 *is unstable to shape deformation on the dashed curve, which begins at 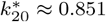. This value is the asymptotic prediction of the pitchfork bifurcation point in Figure 5(a) drawn by a filled star. Parameter Set A of Table 1 was used.*

**Figure 9:**
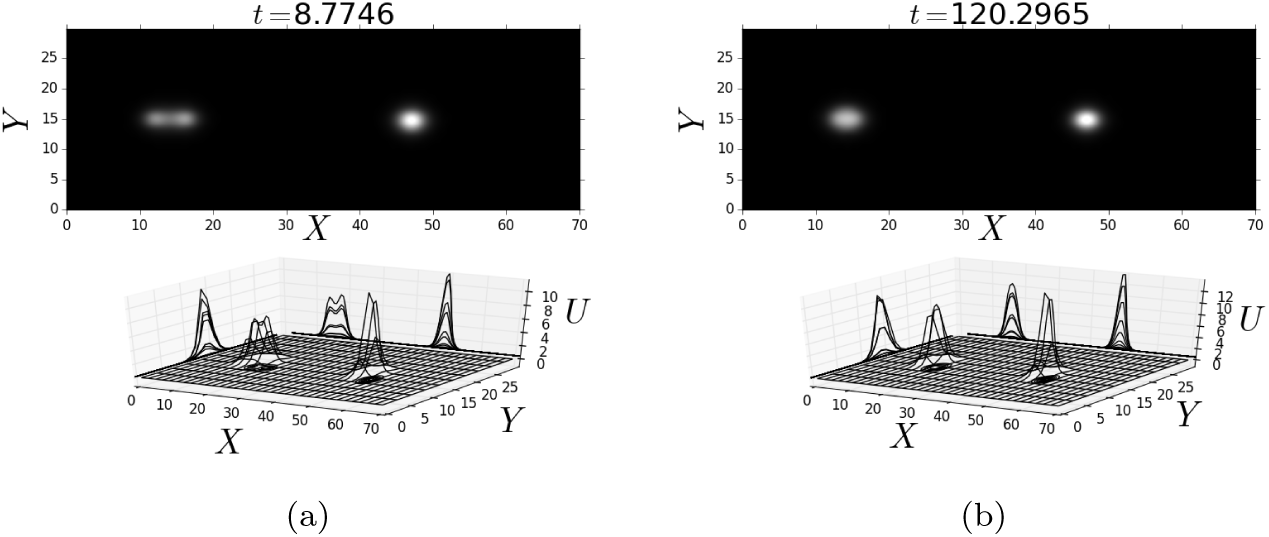
*Two snapshots of a time-dependent numerical simulation where a peanut structure merges into a spot. (a) Peanut structure and a spot. (b) Two spots. Parameter Set A as given in Table 1 and k*_20_ = 0.6723 *for an auxin gradient of the type (i) in* (1.1)

Our main new goal in this paper is to analyze these 2-D spot patterns, which results from the breakup of a stripe, and characterize their slow evolution towards a true steady-state of the RD system. In particular, we will examine how specific forms of the auxin gradient lead to a steady-state spot pattern where the spots are aligned with the direction of the auxin gradient.

**TABLE 1:** *Two parameter sets in the original and dimensionless re-scaled variables. The fundamental units of length and time are µm and sec, and concentration rates are measured by an arbitrary datum (con) per time unit; k*_20_ *is measured by con*^2^*/s, and diffusion coefficients units are µm/s*^2^.

| Parameter Set A |  | Parameter Set B |  |
| --- | --- | --- | --- |
| *Original* | *Re-scaled* | *Original* | *Re-scaled* |
| $D_1 = 0.075$ | $\varepsilon^2 = 1.02 \times 10^{-4}$ | $D_1 = 0.1$ | $\varepsilon^2 = 3.6 \times 10^{-4}$ |
| $D_2 = 20$ | $D = 0.51$ | $D_2 = 10$ | $D = 0.4$ |
| $k_1 = 0.008$ | $\tau = 18.75$ | $k_1 = 0.01$ | $\tau = 11$ |
| $b = 0.008$ | $\beta = 6.25$ | $b = 0.01$ | $\beta = 1$ |
| $c = 0.1$ | | $c = 0.1$ | |
| $r = 0.05$ | | $r = 0.01$ | |
| $k_{20} \in [10^{-3}, 2.934]$ | $\gamma \in [150, 0.051]$ | $k_{20} \in [0.01, 1.0]$ | $\gamma \in [11, 0.11]$ |
| $L_x = 70$ | | $L_x = 50$ | |
| $L_y = 29.848$ | $d_y = 0.4211$ | $L_y = 20$ | $d_y = 0.40$ |

In the RD model below, *U* (**X**, *T*) and *V* (**X**, *T*) denote active- and inactive-ROP densities, respectively, at time *T >* 0 and point 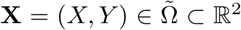, where 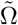, is the rectangular domain 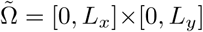. The on-and-off switching interaction is assumed to take place on the cell membrane, which is coupled through a standard Fickian-type diffusive process for both densities (cf. [6, 33]). Although RH cells are flanked by non-RH cells (cf. [17]), there is in general no exchange of ROPs between them. Therefore, no-flux boundary conditions are assumed on 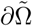. On the other hand, we will consider two specific forms for the spatially-dependent dimensionless auxin distribution *α*: (i) a steady monotone decreasing longitudinal gradient, which is constant in the transverse direction, and (ii) a steady monotone decreasing longitudinal gradient, which decreases on either side of the midline *Y* = *L*_*y*_*/*2; see the left- and right-hand panels in Figure 1, respectively. These two forms are modelled by

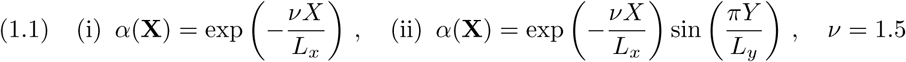

As formulated in [4] and [6], the basic dimensionless RD model is

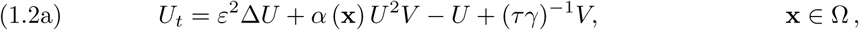

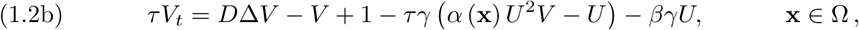

with homogeneous Neumann boundary conditions, and where Ω = [0, 1] *×* [0, *d*_*y*_]. Here Δ is the 2-D Laplacian, and the domain aspect ratio *d*_*y*_ *≡ L*_*y*_*/L*_*x*_ is assumed to satisfy 0 *< d*_*y*_ *<* 1, which represents a cell as is shown in Figure 2. The other dimensionless parameters in (1.2) are defined in terms of the original parameters, as in [4, 6], by

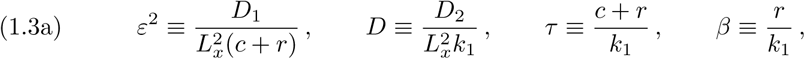

and the primary bifurcation parameter *γ* in this system is given by

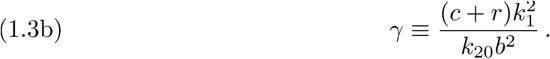

In the original dimensional model of [6], *D*_1_ ≪ *D*_2_ are the diffusivities for *U* and *V, b* is the rate of production of inactive ROP (source parameter), *c* is the rate constant for deactivation, *r* is the rate constant describing active ROPs being used up in cell softening and subsequent hair formation (loss parameter), and the activation step is assumed to be proportional to *k*_1_*V* + *k*_20_*α*(**x**)*U* ^2^*V*. The activation and overall auxin level within the cell, initiating an autocatalytic reaction, is represented by *k*_1_ *>* 0 and *k*_20_ *>* 0 respectively. The parameter *k*_20_, arising in the bifurcation parameter *γ* of (1.3b), will play a key role for pattern formation (see [6, 33] for more details on the model formulation).

We will examine the role that geometrical features, such as the 2-D domain and the auxin-gradient, play on the dynamics of localized regions, referred to as spots or patches, where the active-ROP has an elevated concentration. This analysis is an extension to the analysis performed in [4], where it was shown that a 1-D localized stripe pattern of active-ROP in a 2-D domain will, generically, exhibit breakup into spatially localized spots. To analyze the subsequent dynamics and stability of these localized spots in the presence of the auxin gradient, we will extend the hybrid asymptotic-numerical methodology developed in [25, 35] for prototypical RD systems with spatially homogeneous coefficients. This analysis will lead to a novel finite-dimensional dynamical system characterizing slow spot evolution. Although in our numerical simulations we will focus on the two specific forms for the auxin gradient in (1.1), our analysis applies to an arbitrary gradient.

The outline of the paper is as follows. In section 2 we introduce the basic scaling for (1.2), and we use singular perturbation methods to construct an *N*-spot quasi steady-state solution. In section 3 we derive a differential algebraic system (DAE) of ODEs coupled to a nonlinear algebraic system, which describes the slow dynamics of the centers of a collection of localized spots. In particular, we explore the role that the auxin gradient (i) in (1.1) has on the ultimate spatial alignment of the localized spots that result from the breakup of an interior stripe. For the auxin gradient (i) of (1.1), in section 4 we study the onset of spot self-replication instabilities and other 𝒪(1) time-scale instabilities of quasi steady-state solutions. In section 5 we examine localized spot patterns for the more biologically realistic model auxin (ii) of (1.1). To illustrate the theory, throughout the paper we perform various numerical simulations and numerical bifurcation analyses using the parameter sets in Table 1, which are qualitatively close to the biological parameters (see [6, 33]). Finally, a brief discussion is given in section 6.

## 2. Asymptotic Regime for Spot Formation

We shall assume a shallow, oblong cell, which will be modelled as a 2-D flat rectangular domain, as shown in Figure 2. In this domain, the biochemical interactions leading to an RH initiation process are assumed to be governed by the RD model (1.2).

For a RD system, a spatial pattern in 2-D consisting of spots is understood as a collection of structures where, at least, one of its solution components is highly localized. These structures typically evolve in such a manner that the spatial locations of the spots change slowly in time. However, depending on the parameter regime, these localized patterns can also exhibit fast 𝒪(1) time-scale instabilities, leading either to spot creation or destruction. For prototypical RD systems such as the Gierer–Meinhardt, Gray–Scott, Schnakenberg, and Brusselator models, the slow dynamics of localized solutions and the possibility of self-replication and competition instabilities, leading either to spot creation or destruction, respectively, have been studied using a hybrid asymptotic-numerical approach in [26, 35]. In addition, in the large inhibitor diffusion limit, a leading-order-in--1*/* log *ε* analysis, shows that the linear stability problem for localized spot patterns characterizing competition instabilities can be reduced to the study of classes of nonlocal eigenvalue problems (NLEPs) (cf. [37, 38]). In a 1-D context, rigorous results for the spectral properties of “far-from-equilibrium” periodic 1-D patterns have been obtained recently in [11].

Our previous studies in [6] of 1-D spike patterns and in [4] of 1-D stripe patterns have extended these previous NLEP analyses of prototypical RD systems to the more complicated, and biologically realistic, plant root-hair system (1.2). In the analysis below we extend the 2-D hybrid asymptotic-numerical methodology developed in [26, 35] for prototypical RD systems to (1.2). The primary new feature in (1.2), not considered in these previous works, is to analyze the effect of a spatial gradient in the right-hand sides of (1.2), which represents variations in the auxin distribution. This spatial gradient is central to the spatial alignment of the localized structures.

To initiate our hybrid approach for (1.2), we first need to rescale the RD model (1.2) in Ω in order to determine the parameter regime where localized spots exist. To do so, we integrate the steady-state version of the RD model (1.2) over the domain to show, as in Proposition 2.2 of [6] for the 1-D problem, that for any steady-state solution *U*_0_(**x**) of (1.2) we must have

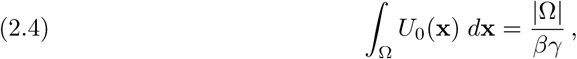

where *|*Ω*|* = *d*_*y*_ is the area of Ω. Since the parameters *d*_*y*_, *β* and *γ* are 𝒪 (1), the constraint (2.4) implies that the average of *U* across the whole domain is also 𝒪 (1). In other words, if we seek localized solutions such that *U →*0 away from localized *O*(*ε*) regions near a collection of spots, then from (2.4) we must have that *U* = *𝒪* (*ε*^−2^) near each spot.

### 2.1 A Multiple-Spot Quasi Steady-State Pattern

In order to derive a finite-dimensional dynamical system for the slow dynamics of active-ROP spots, we must first construct a quasi steady-state pattern describing ROP aggregation in *N* small distinct spatial regions.

To do so, we seek a quasi steady-state solution where *U* is localized within an 𝒪 (*ε*) region near each spot location **x**_*j*_ for *j* = 1, *…, N*. From the conservation law (2.4) we get that *U* = 𝒪 *(ε*^−2^) and, as a consequence, *V* = 𝒪 (*ε*^2^) near each spot. In this way, if we replace *U* = *ε*^−2^*U*_*j*_ near each spot, and define ***ξ*** = *ε*^−1^ (**x** − **x**_*j*_), we obtain from (2.4) that 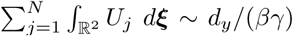. This scaling law motivates the introduction of the new variables *u* and *v* defined by

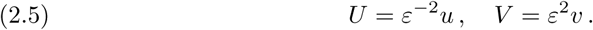

In terms of these new variables, (1.2) with *∂*_*n*_*u* = *∂*_*n*_*v* = 0 on **x** *∈ ∂*Ω becomes

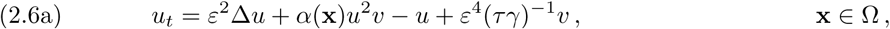

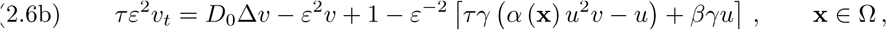

where *D* = *ε*^−2^*D*_0_ comes from balancing terms in (2.6b).

In the inner region near the *j*-th spot, the leading-order terms in the inner expansion are locally radially symmetric. We expand this inner solution as

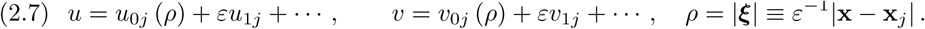

Substituting (2.7) into the steady-state problem for (2.6), we obtain the following leading-order radially symmetric problem on 0 *< ρ < ∞*:

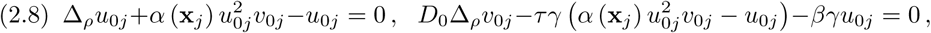

where Δ_*ρ*_ *≡ ∂*_*ρρ*_ + *ρ*^−1^*∂*_*ρ*_ is the Laplacian operator in polar co-ordinates. As a remark, in our derivation of the spot dynamics in section 3 we will need to retain the next term in the Taylor series of the auxin distribution, representing the auxin gradient in the regions where active-ROP is localized. This is given by

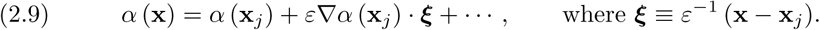

These higher-order terms are key to deriving a dynamical system for spot dynamics. We consider (2.8) together with the boundary conditions

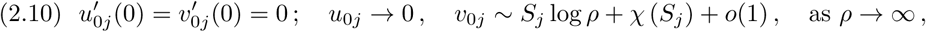

where *S*_*j*_ is called the *source parameter*. The logarithmic far-field behavior for *v*_0*j*_ arises since, owing to the fact that there is no *-v*_0*j*_ term in the second equation of (2.8), we cannot assume that *v*_0*j*_ is bounded as *ρ→∞.* The correction term *X*_*j*_ = *X*(*S*_*j*_) in the far-field behavior is determined from

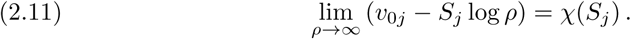

By integrating the equation for *v*_0*j*_ in (2.8), and using the limiting behavior *v*_0*j*_ *∼ S*_*j*_ log *ρ* as *ρ → ∞*, we obtain the identity

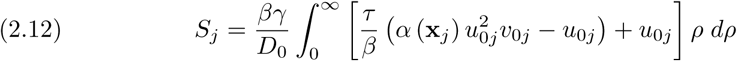

In (2.8), we introduce the change of variables

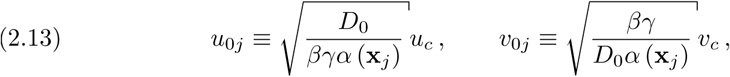

to obtain what we refer to as the *shape canonical core problem* (SCCP), which consists of

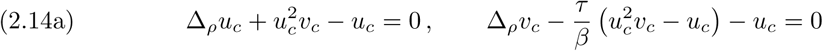

on 0 *< ρ <* ∞, together with the following boundary conditions where *S*_*cj*_ is a parameter:

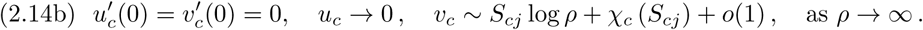

Upon substituting (2.13) into the identity (2.12), we obtain

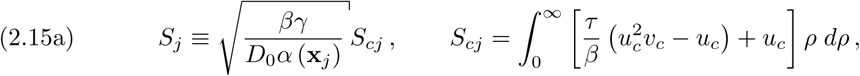

which, from comparing (2.13) and (2.10), yields

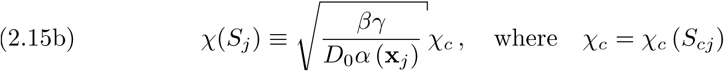

To numerically compute solutions to (2.14) we use the MATLAB code BVP4C and solve the resulting BVP on the large, but finite, interval [0, *ρ*_0_], for a range of values of the parameter *S*_*cj*_. The constant *χ*_*c*_ (*S*_*cj*_) is identified from *v*_*c*_ (*ρ*_0_) *− S*_*c*_ log *ρ*_0_ = *χ*_*c*_, where *v*_*c*_ is the numerically computed solution, and so it depends only on *S*_*cj*_ and the specified ratio *τ/β* in (2.14). We solve the boundary-value problem on 0 *< ρ*_0_ *<* 12 with a tolerance of 10^-4^. Increasing *ρ*_0_ did not change the computational results shown in Figure 3. The results of our computation are shown in Figure 3(a) where we plot *χ*_*c*_ (*S*_*cj*_) versus *S*_*cj*_. In Figure 3(b) we also plot *u*_*c*_ and *v*_*c*_ at a few values of *S*_*cj*_. We observe that *u*_*c*_(*ρ*) has a volcano-shaped profile when *S*_*cj*_ = 5, which suggests, by means of the identity (2.12), that this parameter value should be near where spot self-replication will occur (see subsection 4.1).

We remark that the SCCP (2.14) depends only on the ratio *τ / β*, which characterizes the deactivation and removal rates of active-ROP (see (1.3a)). This implies that the SCCP (2.14) only describes the aggregation process. On the other hand, notice that (2.13) and (2.15) reveal the role that the auxin plays on the activation process. Since *α*(**x**) decreases longitudinally for any of the two forms in (1.1), the scaling (2.13) is such that the further from the left boundary active-ROP localizes, the larger will be the solution amplitude. This feature will be confirmed by numerical simulations in subsection 2.2. In addition, the scaling of the source-parameter in (2.15) indicates that the auxin controls the inactive-ROP distribution along the cell. That is, while *S*_*cj*_ characterizes how *v* interacts with *u* in the region where a spot occurs, the parameter *S*_*j*_ will be determined by the inhomogeneous distribution of source points, which are also governed by the auxin gradient. In other words, the gradient controls and catalyzes the switching process, which leads to localized elevations of concentration of active-ROP.

Next, we examine the outer region away from the spots centered at **x**_1_, *…,* **x**_*N*_, which are locations where *u* is localized. To leading-order, the steady-state problem in (2.6a) yields that *u*_0_ *∼ ε*^4^*v*_0_*/*(*βγ*) as obtained from balancing terms. Therefore, from this order in the reaction terms of (2.6b), we obtain in the outer region that

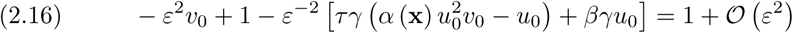

Next, we calculate the effect of the localized spots on the outer equation for *v*. Upon assuming *u ∼ u*_*j*_ and *v ∼ v*_*j*_ near the *j*-th spot, we approximate the singular terms in the sense of distributions as Dirac masses. Indeed, labelling the reactive terms by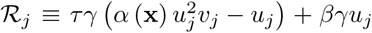, we use (2.12) and the Mean Value Theorem to calculate, in the sense of distributions, that

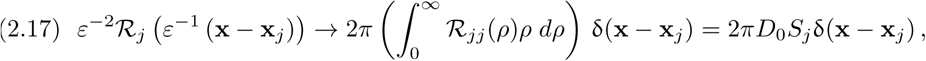

as ε →0, where 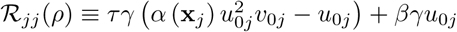.

Combining (2.16), (2.17) and (2.6b) we obtain that the leading-order outer solution *v*_0_, in the region away from *𝒪*(*ε*) neighborhoods of the spots, satisfies

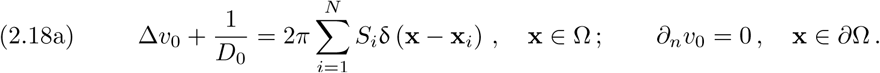

By using (2.10) to match the outer and inner solutions for *v*, we must solve (2.18a) subject to the following singularity behavior as **x** *→* **x**_*j*_:

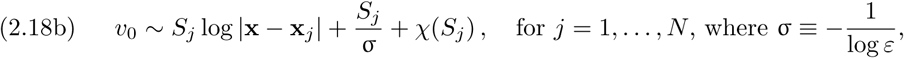

which results from writing (2.10) as *ρ* = *ε*^−1^*|***x** − **x**_*j*_*|*.

We now proceed to solve (2.18) for *v*_0_ and derive a nonlinear algebraic system for the source parameters *S*_*c*1_, *…, S*_*cN*_. To do so, we introduce the unique Neumann Green’s function *G*(**x**; **y**) satisfying (cf. [26])

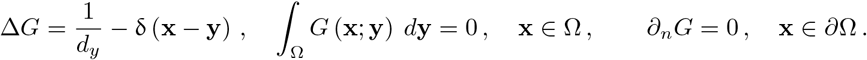

This Green’s function satisfies the reciprocity relation *G* (**x**; **y**) = *G* (**y**; **x**), and can be globally decomposed as *G* (**x**; **y**) = −1*/*(2*π*) log *|***x** − **y***|* + *R* (**x**; **y**), where *R* (**x**; **y**) is the smooth regular part of *G* (**x**; **y**). For **x** *→* **y**, this Green’s function has the local behavior

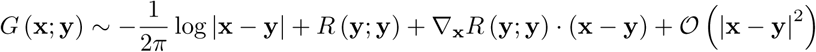

An explicit infinite series solution for *G* and *R* in the rectangle Ω is available (cf. [26]).

In terms of this Green’s function, the solution to (2.18a) is

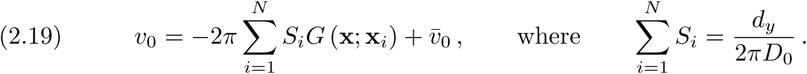

Here 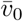 is a constant to be found. The condition on *S*_*i*_ in (2.19) arises from the Divergence Theorem applied to (2.18). We then expand *v*_0_ as **x → x**_*j*_ and enforce the required singularity behavior (2.18b). This yields that

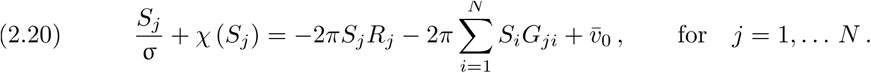

Here *G*_*ji*_ *≡ G* (**x**_*j*_; **x**_*i*_) and *R*_*j*_ *≡ R* (**x**_*j*_; **x**_*j*_). The system (2.20), together with the constraint in (2.19), defines a nonlinear algebraic system for the *N* + 1 unknowns *S*_1_, *…, S*_*N*_ and 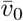 To write this system in matrix form, we define **e** *≡* (1, *…,* 1)^*T*^, **S***≡* (*S*_1_, *…, S*_*N*_)^*T*^, ***χ****≡* (*χ*(*S*_1_), *…, χ*(*S*_*N*_)^*T*^, *E ≡* **ee**^*T*^ */N*, and the symmetric Green’s matrix **G** with entries

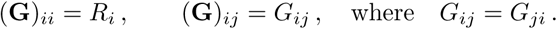

Then, (2.20), together with the constraint in (2.19), can be written as

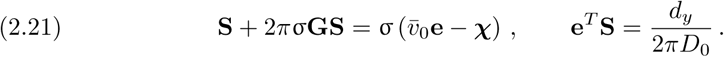

Left multiplying the first equation in (2.21) by **e**^*T*^, and using the second equation in (2.21), we find an expression for 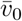

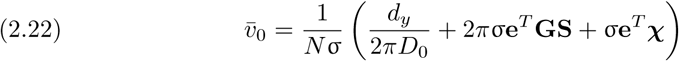

Substituting 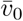 back into the first equation of (2.21), we obtain a nonlinear system for *S*_1_, *…, S*_*N*_ given by

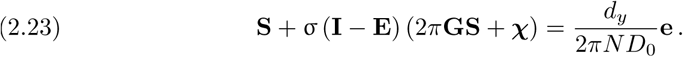

In terms of **S**, 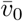 is given in (2.22). To relate *S*_*j*_ and *χ*_*j*_ with the SCCP (2.14), we define

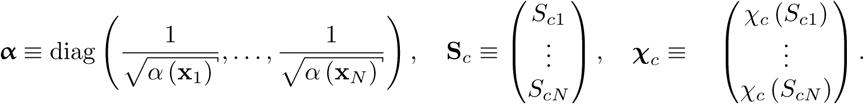

Then, by using (2.15), we find (2.24)

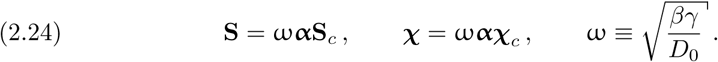

Finally, upon substituting (2.24) into (2.23) we obtain our main result regarding quasi steady-state spot patterns.

PROPOSITION 1. *Let* 0 *< ε ≪* 1 *and assume D*_0_ = 𝒪 (1) *in* (2.6) *so that D =* 𝒪*(ε−*^*2*^*) in* (1.2). *Suppose that the nonlinear algebraic system for the source parameters, given by*

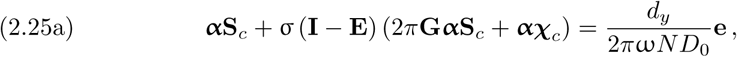

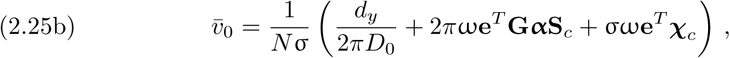

*has a solution for the given spatial configuration* **x**_1_, *…,* **x**_*N*_ *of spots. Then, there is a quasi steady-state solution for* (1.2) *with U* = *ε*^−2^*u and V* = *ε*^2^*v. In the outer region away from the spots, we have u* = 𝒪 (*ε*^4^) *and v ∼ v*_0_, *where v*_0_ *is given asymptotically in terms of* **S** = ω**αS**_*c*_ *by* (2.19). *In contrast, in the j-th inner region near* **x**_*j*_, *we have*

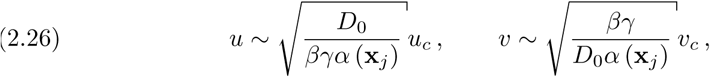

*where u*_*c*_ *and v*_*c*_ *is the solution of the SCCP* (2.14) *in terms of S*_*cj*_ *satisfying* (2.25a).

### 2.2 Spots and Baby Droplets

Proposition 1 provides an asymptotic characterization of a *N*-spot quasi steady-state solution. As has been previously studied in the previous paper [4], localized spots can be generated by the breakup of a straight homoclinic stripe. The numerical simulations in Figure 4, for the 1-D auxin gradient in (1.1), illustrate how small spots are formed from the breakup of an interior homoclinic stripe and how they evolve slowly in time to a steady-state two-spot pattern aligned with the direction of the auxin gradient. The initial condition for these simulations is obtained by trivially extending a 1-D spike in the transversal direction to obtain a homoclinic stripe, which is a steady-state of (1.2), and then perturbing it with a transverse perturbation of small amplitude. As shown in Figure 4(c)–Figure 4(d), the amplitude of the left-most spot decreases as the spot drifts towards the left domain boundary at *X* = 0, while the amplitude of the right-most spot grows as it drifts towards the right domain boundary at *X* = *L*_*x*_. The spot profiles are described by Proposition 1 for each fixed re-scaled variables x_*j*_. Our realizations of the asymptotic theory shown below in Figure 7 confirm the numerical results in Figure 4(d) that the spots become aligned with the direction of the auxin gradient.

In order to explore solutions that bifurcate from a multi-spot steady state, we adapted the MATLAB code of [2, 34] and performed a numerical continuation for the RD system (1.2) in the bifurcation parameter *k*_20_, starting from the solution in Figure 4(d) (all other parameters as in Parameter Set A in Table 1). The results are shown in Figure 5. In the bifurcation diagram depicted in Figure 5(a) the linearly stable branch is plotted as a solid curve and unstable ones by light gray-dashed curves. We found fold bifurcations and a pitchfork bifurcation, represented by a filled circle and a filled star, respectively. To the right of the pitchfork bifurcation, the symmetric solution branch destabilises; the emerging branch of asymmetric states is fully unstable, and features a sequence of fold bifurcations. Overall, these results suggest that a snaking bifurcation structure occurs in 2-D domains, similar to what was found in [4] for the corresponding 1-D system. This intricate branch features a set of fold points (not explicitly labelled in Figure 5(a)), and around each of them a new spot emerges. A pattern consisting of a spot and a peanut structure is found in the solution branch labelled as P. This solution is shown in Figure 5(c).

We found that solutions with peanut structures or baby droplets are unstable steady-states in this region of parameter space, and so they do not persist in time; see a sample of these in Figure 5(d). In addition, as can be seen in Figure 5(b), there are stable multi-spot solutions when *k*_20_ is small enough. Overall, this suggests that certain 𝒪(1) time-scale instabilities play an important role in transitioning between steady-state patterns (cf. [8, 20]). Further, in section 4, peanut-splitting instabilities are addressed.

## 3. The Slow Dynamics of Spots

In subsection 2.1 we constructed an *N*- spot quasi steady-state solution for an arbitrary, but “frozen”, spatial configuration **x**_1_, *…,* **x**_*N*_ of localized patches of active-ROP. In this section, we derive a differential-algebraic system of ODEs (DAE) for the slow time-evolution of **x**_1_, *…,* **x**_*N*_ towards their equilibrium locations in Ω. This system will consist of a constraint that involves the slow time evolution of the source parameters *S*_*c*1_, *…, S*_*cN*_. The derivation of this DAE system is an extension of the 1-D analysis of [6].

To derive the DAE system, we must extend the analysis in subsection 2.1 by retaining the 𝒪(*ε*) terms in (2.7) and (2.9). By allowing the spot locations to depend on a distinguished limit for the slow time scale *η* = *ε*^2^*t* as **x**_*j*_ = **x**_*j*_ (*η*), we obtain in the *j*-th inner region that the corrections terms *u*_1*j*_ and *v*_1*j*_ to the SCCP (2.14) satisfy

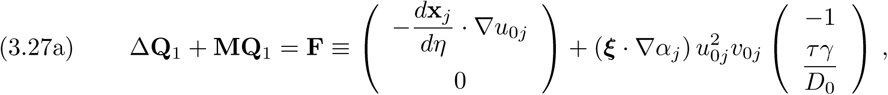

where the vector **Q**_1_ and the matrix **M** are defind by

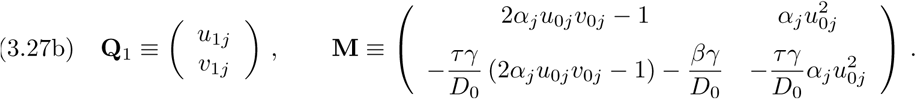

Here *α*_*j*_ *≡ α* (**x**_*j*_) and *∇α*_*j*_ *≡ ∇α*(**x**_*j*_). To determine the far-field behavior for **Q**_1_, we expand the outer solution *v*_0_ in (2.19) as **x** *→* **x**_*j*_ and retain the 𝒪 (*ε*) terms. This yields

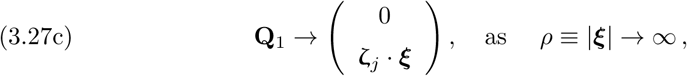

where, in terms of the source strengths *S*_*j*_ defined from (2.23), we have

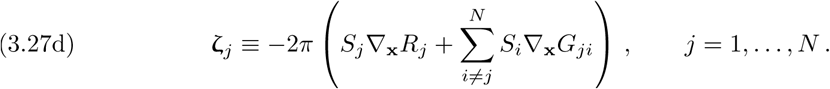

Here *∇***_x_***R*_*j*_ *≡* _**x**_*R* (**x**_*j*_; **x**_*j*_) and *∇***_x_** *G*_*j*_ *≡ ∇***_x_** *G* (**x**_*j*_; **x**_*j*_) for *j* = 1, *…, N*.

To impose a solvability condition on (3.27), which will lead to ODEs for the spot locations, we first rewrite (3.27) in terms of the canonical variables (2.13) of the SCCP (2.14). Using this transformation, the right-hand side **F** of (3.27a) becomes

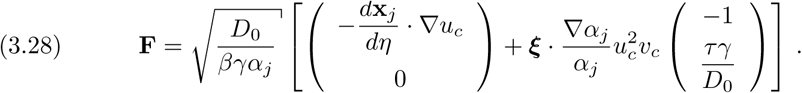

In addition, the matrix **M** on the left-hand side of (3.27a) becomes

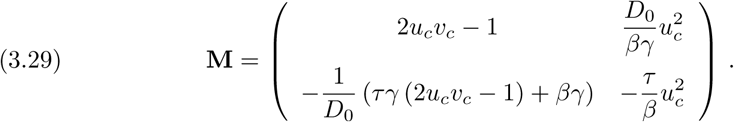

Together with (3.28) this suggests that we define new variables 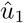and 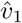 by

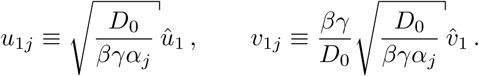

In terms of these new variables, and upon substituting (3.28) and (3.29) into (3.27), we obtain that 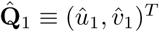 in ξ ∈ ℝ^2^ satisfies

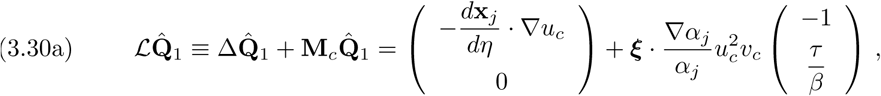

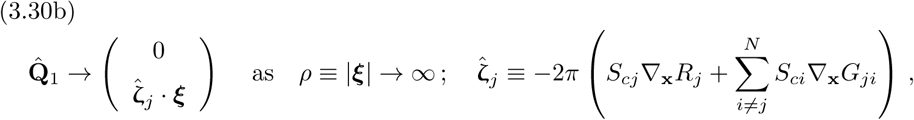

and *S*_*cj*_ for *j* = 1, *…, N* satisfies the nonlinear algebraic system (2.25a). In addition, in (3.30a) **M**_*c*_ is defined by

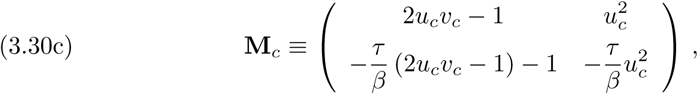

where *u*_*c*_ and *v*_*c*_ satisfy the SCCP (2.14), which is parametrized by *S*_*cj*_.

In (3.30b), the source parameters *S*_*cj*_ depend indirectly on the auxin distribution through the nonlinear algebraic system (2.25), as summarized in Proposition 1. We further observe from the second term on the right-hand side of (3.30a) that its coefficient *∇α*_*j*_*/α*_*j*_ is independent of the magnitude of the spatially inhomogeneous distribution of auxin, but instead depends on the direction of the gradient.

### 3.1 Solvability Condition

We now derive a DAE system for the spot dynamics by applying a solvability condition to (3.30). We label ***ξ*** = (*ξ*_1_, *ξ*_2_)^*T*^ and **u**_*c*_ *≡* (*u*_*c*_, *v*_*c*_)^*T*^. Differentiating the core problem (2.14a) with respect to the *ξ*_*i*_, we get

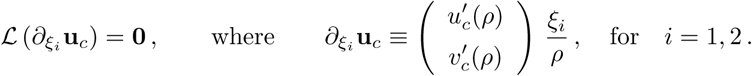

This demonstrates that the dimension of the nullspace of *ℒ* in (3.30), and hence its adjoint *ℒ\**, is at least two-dimensional. Numerically, for our two parameter sets in Table 1, we have checked that the nullspace of *ℒ* is exactly two-dimensional, provided that *S*_*j*_ does not coincide with the spot self-replication threshold given in subsection 4.1.

There are two independent nontrivial solutions to the homogeneous adjoint problem 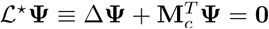 given by Ψ_*i*_ ≡ **P**(ρ) ξ_*i*_/ρ for *i* = 1,2 where **P** satisfies

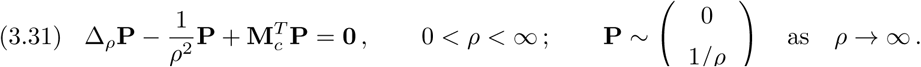

Here the condition at infinity in (3.31) is used as a normalization condition for **P**.

Next, to derive our solvability condition we use Green’s identity over a large disk Ω_*ρ*0_ 1 of radius |***ξ|*** = *ρ*_0_ ≫ to obtain for *i* = 1, 2 that

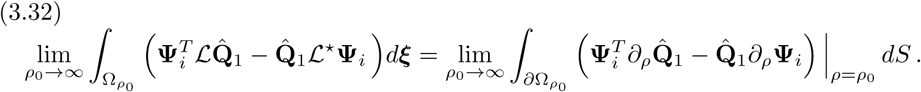

With **Ψ**_*i*_ *≡* **P**(*ρ*)*y*_*i*_*/ρ*, for *i ∈ {*1, 2*}*, we first calculate the left-hand side (LHS) of (3.32) using (3.30a) to obtain **P** ≡ (*P*_1_(ρ), *P*_1_(ρ))^*T*^ with 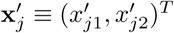 that

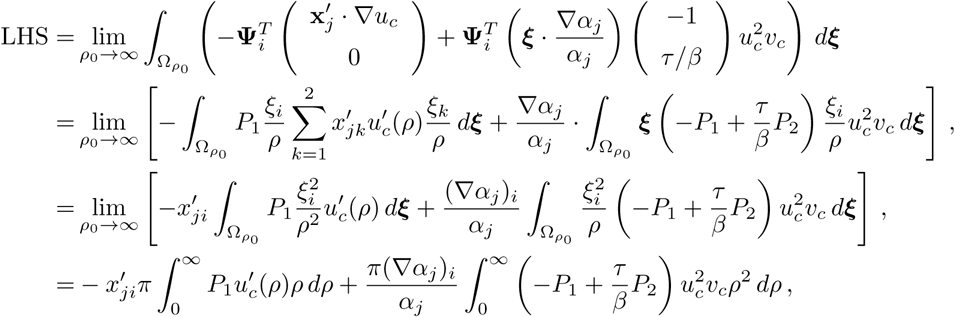

where 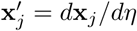. In deriving the result above, we used the identity 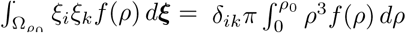 for any radially symmetric function f(*ρ*),where δ_ik_ is the Kronecker delta.

Next, we calculate the right-hand side (RHS) of (3.32) using the far-field behaviors of 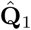 and **P** as *ρ → ∞*. We derive that

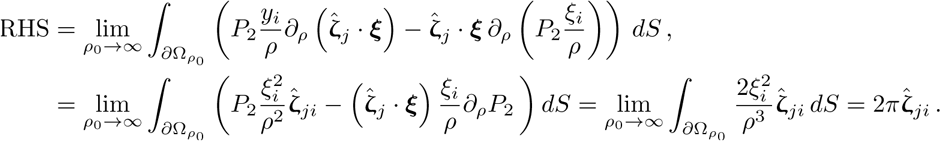

In the last passage, we used *dS* = *ρ*_0_*dθ* where *θ* is the polar angle. Finally, we equate LHS and RHS for *i* = 1, 2, and write the resulting expression in vector form:

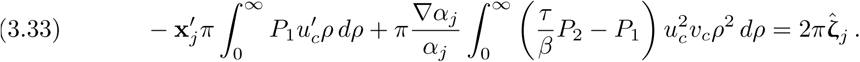

We summarize our main, formally-derived, asymptotic result for slow spot dynamics as follows:

PROPOSITION 2. *Under the same assumptions as Proposition 1, and assuming that the N-spot quasi steady-state solution is stable on an* 𝒪(1) *time-scale, the slow dynamics on the long time-scale η* = *ε*^2^*t of this quasi steady-state spot pattern consists of the constraints* (2.25) *coupled to the following ODEs for j* = 1, *…, N:*

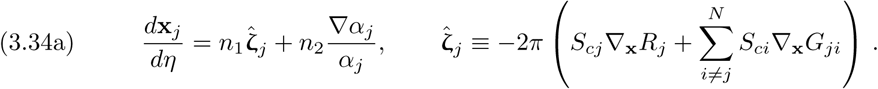

*The constants n*_1_ *and n*_2_, *which depend on S*_*cj*_ *and the ratio τ/β are defined in terms of the solution to the SCCP* (2.14) *and the homogeneous adjoint solution* (3.31) *by*

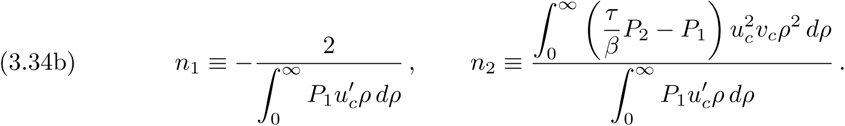

*In* (3.34a), *the source parameters S*_*cj*_ *satisfy the nonlinear algebraic system* (2.25), *which depends on the instantaneous spatial configuration of the N spots. Overall, this leads to a DAE system characterizing slow spot dynamics.*

PROPOSITION 2 describes the slow dynamics of a collection of *N* localized spots under an arbitrary, but smooth, spatially-dependent auxin gradient. It is an extension of the 1-D analysis of spike evolution, considered in [6]. The dynamics in (3.34a), shows that the spot locations depend on the gradient of the Green’s function, which depends on the domain Ω, as well as the spatial gradient of the auxin distribution. In particular, the spot dynamics depends only indirectly on the magnitude of the auxin distribution *α*(**x**_*j*_) through the source parameter *S*_*cj*_. The auxin gradient *∇α*_*j*_, however, is essential to determining the true steady-state spatial configuration of spots. In addition, the spatial interaction between the spots arises from the terms in the finite sum of (3.34a), mediated by the Green’s function. Since the Green’s function and its regular part can be found analytically for a rectangular domain (cf. [26]), we can readily use (3.34) to numerically track the slow time-evolution of a collection of spots for a specified auxin gradient.

Before illustrating results from the DAE dynamics, we must determine *n*_1_ and *n*_2_ as a function of *S*_*cj*_ for a prescribed ratio *τ/β*. This ratio is associated with the linear terms in the kinetics of the original dimensional system (1.2), which are related to the deactivation of ROPs and production of other biochemical complexes which promote cell wall softening (cf. [6, 33]). To determine *n*_1_ and *n*_2_, we first solve the adjoint problem (3.31) numerically using the MATLAB routine BVP4C. This is done by enforcing the local behavior that **P** = 𝒪 (*ρ*) as *ρ →* 0 and by imposing the far-field behavior for **P**, given in (3.31), at *ρ* = *ρ*_0_ = 12. In Figure 6(a) we plot *P*_1_ and *P*_2_ for three values of *S*_*cj*_, where *τ/β* = 3, and Parameter Set A in Table 1 was used. For each of the three values of *S*_*cj*_, we observe that the far-field behavior *P*_2_(*ρ*) *∼* 1*/ρ* and *P*_1_(*ρ*) *∼* (*τ/β -* 1) */ρ* as *ρ → ∞*, which is readily derived from (3.31), is indeed satisfied. Upon performing the required quadratures in (3.34b), in Figure 6(b) we plot *n*_1_ and *n*_2_ versus *S*_*cj*_. These numerical results show that *n*_1_ *>* 0 and *n*_2_ *>* 0 for *S*_*cj*_ *>* 0, which will ensure existence of stable fixed points of the DAE dynamics. These stable fixed points correspond to realizable steady-state spot configurations for our two specific forms for the auxin gradient.

### 3.2 Comparison Between Theory and Asymptotics for Slow Spot Dynamics

In this subsection we compare predictions from our asymptotic theory for slow spot dynamics with corresponding full numerical results computed from (2.6) using a spatial mesh with 500 and 140 gridpoints in the *x* and *y* directions, respectively. For the time-stepping a modified Rosenbrock formula of order 2 was used.

The procedure to obtain numerical results from our asymptotic DAE system in Proposition 2 is as follows. This DAE system is solved numerically by using Newton’s method to solve the nonlinear algebraic system (2.25) together with a Runge–Kutta ODE solver to evolve the dynamics in (3.34a). The solvability integrals *n*_1_(*S*_*cj*_) and *n*_2_(*S*_*cj*_) in the dynamics (3.34a) and the function χ_c_ in the nonlinear algebraic system (2.25) are pre-computed at 200 grid points in *S*_*cj*_ and a cubic spline interpolation is fitted to the discretely sampled functions to compute them at arbitrary values of *S*_*cj*_. For the rectangular domain, explicit expressions for the Green’s functions *G* and *R*, together with their gradients, as required in the DAE system, are calculated from the expressions in §4 of [26].

To compare our results for slow spot dynamics we use Parameter Set A of Table 1 and take *k*_20_ = 0.3. For the auxin gradient we took the monotone 1-D gradient in type (i) in (1.1). By a small perturbation of the unstable 1-D stripe solution, our full numerical computations of (2.6) lead to the creation of two localized spots. Numerical values for the centers of the two spots are calculated (see the caption of Figure 7) and these values are used as the initial conditions for our numerical solution of the DAE system in Proposition 2. In Figure 7 we compare our full numerical results for the *x* and *y* coordinates for the spot trajectories, as computed from the RD system (2.6), and those computed from the corresponding DAE system. The key distinguishing feature in these results is that the spots become aligned to the longitudinal midline of the domain. We observe that the *y*-components of the spot trajectories are predicted rather accurately over long time intervals by the DAE system. However, although the *x*-components of the trajectories are initially close, they deviate somewhat as *t* increases. Since we only have a formal asymptotic theory, with no error bounds, and because it is computationally expensive to perform more refined numerical simulations over very-long time scales from the full RD system, we are not able to precisely identify why the agreement in the *y*-coordinate is better than for the *x*-coordinate. However, overall, the asymptotic theory accurately identifies the time-scale over which the two localized spots become aligned with the auxin gradient along the mid-line of the cell.

## 4. Fast 𝒪(1) Time-Scale Instabilities

We now briefly examine the stability properties of the *N*-spot quasi-equilibrium solution of Proposition 1 to 𝒪(1) time-scale instabilities, which are fast relative to the slow dynamics. We first consider spot self-replication instabilities associated with non-radially symmetric perturbations near each spot.

### 4.1 The Self-Replication Threshold

Since the speed of the slow spot drift is 𝒪 (*ε*^2^) *≪* 1, in our stability analysis below we will assume that the spots are “frozen” at some configuration **x**_1_, *…,* **x**_*N*_. We will consider the possibility of instabilities that are locally non-radially symmetric near each spot. In the inner region near the *j*-th spot at **x**_*j*_, where ***ξ*** = *ε*^−1^(**x–x**_*j*_), we linearize (2.6) around the leading-order core solution *u*_0*j*_, *v*_0*j*_, satisfying (2.8), by writing 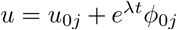 and *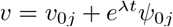*. From (2.6), we obtain to leading order that

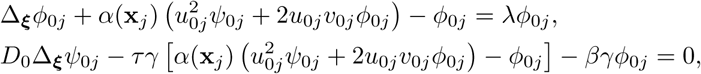

where Δ**_*ξ*_** is the Laplacian in the local variable ***ξ***. Then, upon relating *u*_0*j*_, *v*_0*j*_ to the SCCP by using (2.13), and defining 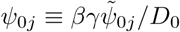 the system above reduces to

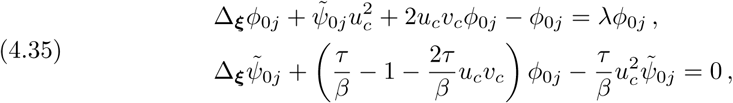

where *u*_*c*_, *v*_*c*_ is the solution to the SCCP (2.14).

We then look for an 𝒪 (1) time-scale instability associated with the local angular integer mode *m ≥* 1 by introducing the new variables Φ_0_(*ρ*) and Ψ_0_(*ρ*) defined by

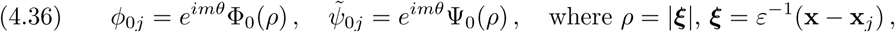

and ***ξ***^*T*^ = *ρ*(cos *θ,* sin *θ*). Substituting (4.36) into (4.35), we obtain the eigenvalue problem:

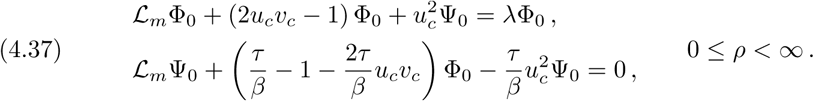

Here we have defined *ℒ*_*m*_γ *≡ ∂*_*ρρ*_γ+*ρ*^−1^*∂*_*ρ*_γ*-m*^2^*ρ*^−2^γ. We impose the usual regularity condition for Φ_0_ and Ψ_0_ at *ρ* = 0. The appropriate far-field boundary conditions for (4.37) is discussed below.

Since the eigenvalue problem (4.37) is difficult to study analytically, we solve it numerically for various integer values of *m*. We denote *λ*_max_ to be the eigenvalue of (4.37) with the largest real part. Since *u*_*c*_ and *v*_*c*_ depend on *S*_*cj*_ from the SCCP (2.14), we have implicitly that *λ*_max_ = *λ*_max_(*S*_*cj*_, *m*). To determine the onset of any instabilities, for each *m* we compute the smallest threshold value *S*_*cj*_ *= ∑* _*m*_ where Re(λ_*max*_(*∑*_*m*_,*m*)) =0.

In our computations, we only consider *m* = 2, 3, 4, *…*, since *λ*_max_ = 0 for any value of *S*_*cj*_ for the translational mode *m* = 1. Any such instability for *m* = 1 is reflected in instability in the DAE system (3.34a).

For *m ≥* 2 we impose the far-field behavior that Φ_0_ decays exponentially as *ρ*_*0*_ *→ ∞* while Ψ_0_ *∼ 𝒪*(*ρ*^*-m*^) *→* 0 as *ρ → ∞*. With this limiting behavior, (4.37) is discretized with centered differences on a large but finite domain. We then determine *λ*_max_(*S*_*cj*_, *m*) by computing the eigenvalues of the discretized eigenvalue problem, in matrix form. For *m ≥* 2 our computations show that *λ*_max_(*S*_*cj*_, *m*) is real and that *λ*_max_(*S*_*cj*_, *m*) *>* 0 if and only if S_cj_ > Σ_m_. In our computations we took 400 meshpoints on the interval 0 *≤ ρ <* 15. For the ratio *τ/β* = 3, corresponding to Parameter Set A of Table 1, the results for the threshold values Σ_m_ for *m* = 2, 3, 4 given in Figure 8(a) are insensitive to increasing either the domain length or the number of grid points. Our main conclusion is that as *S*_*cj*_ is increased, the solution profile of the *j*-th spot first becomes unstable to a non-radially symmetric peanut-splitting mode, corresponding to *m* = 2, as *S*_*cj*_ increases above the threshold ∑_2_ ≈ 4.16 when *τ/β* = 3. As a remark, if we take *τ/β* = 11, corresponding to Parameter Set B of Table 1, we compute instead that Σ_2_ ≈ 3.96.

To illustrate this peanut-splitting threshold, we consider a pattern with a single localized spot. Using (2.19) we find *S*_1_ = *d*_*y*_*/*(2*πD*_0_). Then, (2.15) yields the following expression for the source parameter for the SCCP (2.14)

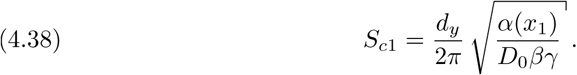

When *τ/β* = 3, corresponding to Parameter Set A of Table 1, we predict that the one-spot pattern will first undergo a shape-deformation, due to the *m* = 2 peanut-splitting mode, when *S*_*c*1_ exceeds the threshold ∑_2_ ≈ 4.16. Since *γ* is inversely proportional to the parameter *k*_20_, as shown in (1.3b), which determines the auxin level, we predict that as the auxin level is increased above a threshold a peanut structure will develop from a solitary spot. As shown below in subsection 4.2 for an auxin distribution of the type (i) in (1.1), this linear peanut-splitting instability triggers a nonlinear spot replication event.

*In addition, in Figure 8(b) we plot the source parameters corresponding to the two-spot steady-state aligned along the midline Y* = *L*_*y*_*/*2 as the parameter *k*_20_ is varied. These asymptotic results, which asymptotically characterize states in branch A of Figure 5(a), are computed from the steady-state of the DAE system (3.34). Notice that the spot closest to the left-hand boundary loses stability at 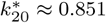 when *S*_*c*1_ meets the peanut-splitting threshold ∑_2_. On the other hand, *S*_*c*2_ values corresponding to the spot closest to the right-hand boundary are always below Σ_2_ for the parameter range considered. Hence, no shape change for this spot is observed. This result rather accurately predicts the pitchfork bifurcation in Figure 5(a) portrayed by a filled star at 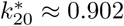.

### 4.2 Numerical Illustrations of 𝒪 (1) Time-Scale Instabilities

In Figure 5(c) we showed a steady-state solution consisting of a spot and a peanut structure, which is obtained from numerical continuation in the parameter *k*_20_. This solution belongs to the unstable branch labelled by P in Figure 5(a), which is confined between two fold bifurcations (not explicitly shown). We take a solution from this branch as an initial condition in Figure 9(a) for a time-dependent numerical simulation of the full RD model (1.2)-(2.6) using a centered finite-difference scheme. Our numerical results in Figure 9(b) show that this initial state evolves dynamically into a stable two-spot solution. This is a result of the overlapping of the stable branch A with an unstable one in the bifurcation diagram.

Other initial conditions and parameter ranges also exhibit 𝒪 (1) time-scale instabilities. To illustrate these instabilities, and see whether a self-replication spot process is triggered by a linear peanut-splitting instability, we perform a direct numerical simulation of the full RD model (1.2) taking as initial condition the unstable steady-state solution labelled by B in Figure 5(a). This steady-state, shown in Figure 10(a), consists of two spots, with one having four small droplets associated with it. This particular steady-state is chosen since both competition and self-replication instabilities can be seen in the time evolution of this initial condition in the panels of Figure 10. Firstly, we observe in Figure 10(b) that the small droplets for the left-most spot in Figure 10(a) are annihilated in 𝒪 (1) times. Then, the right-most spot, which has a relatively low source parameter owing to its distance away from the left-hand boundary (see (2.15a)), a source parameter *S*_*c*1_ = 9.1403, given in (4.38) of subsection 4.1, that exceeds the peanut-splitting threshold ∑_2_ ≈ 4.16. This linear shape-deforming instability gives rise to the peanut structure shown in Figure 10(d), and the direction of spot-splitting is aligned with the direction of the monotone decreasing auxin gradient, which depends only on the longitudinal direction. In Figure 10(e) and Figure 10(f) we show that this linear peanut-splitting instability has triggered a nonlinear spot-splitting event that creates a second localized spot. Finally, these two spots drift away from each other and their resulting slow dynamics is characterized by the DAE dynamics in Proposition 2.

**Figure 10:**
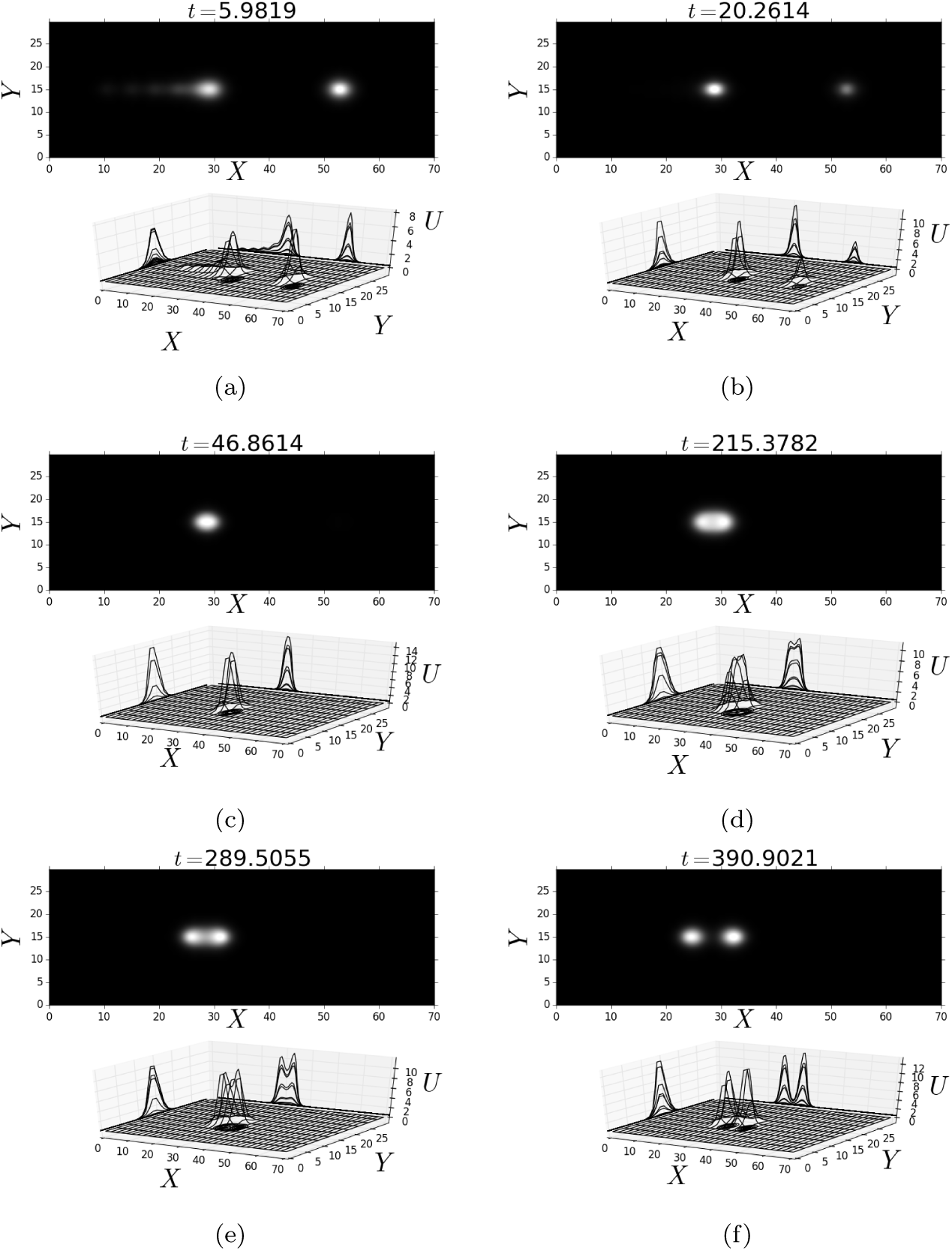
*Numerical simulations of the full RD system illustrating* 𝒪(1) *time-scale instabilities. Competition instability: (a) baby droplets and (b) a spot gets annihilated. Spot self-replication: (c) one spot, (d) early stage of a self-replication process, (e) a clearly visible peanut structure, and (f) two distinct spots moving away from each other. Parameter Set A as given in Table 1 and k*_20_ = 1.6133.

Competition and self-replication instabilities, such as illustrated above, are two types of fast 𝒪(1) time-scale instabilities that commonly occur for localized spot patterns in singularly perturbed RD systems. Although it is beyond the scope of this paper to give a detailed analysis of a competition instability for our plant RD model (1.2), based on analogies with other RD models (cf. [9, 25, 26, 35]), this instability typically occurs when the source parameter of a particular spot is below some threshold or, equivalently, when there are too many spots that can be supported in the domain for the given substrate diffusivity. In essence, a competition instability is an overcrowding instability. Alternatively, as we have unravelled in subsection 4.1, the self-replication instability is an undercrowding instability and is triggered when the source parameter of a particular spot exceeds a threshold, or equivalently when there are too few spots in the domain. For the standard Schnakenberg model with spatially homogeneous coefficients, in [26] it was shown through a center-manifold type calculation that the direction of splitting of a spot is always perpendicular to the direction of its motion. However, with an auxin gradient the direction of spot-splitting is no longer always perpendicular to the direction of motion of the spot. In our plant RD model, the auxin gradient not only enhances robustness of solutions, which results in the overlapping of solution branches and is suggestive that a strong slanted homoclinic snaking mechanism can occur (see the next section), but it also controls the location of steady-state spots (see subsection 2.2 and section 3).

## 5. More Robust 2D Patches and Auxin Transport

We now consider a more biologically plausible model for the auxin transport, which initiates the localization of ROP. This model allows for both a longitudinal and transverse spatial dependence of auxin. The original ROP model, derived in [6, 33] and analyzed in [4, 6] and the sections above, depends crucially on the spatial gradient of auxin. The key assumption we have made above was to assume a decreasing auxin distribution along the *X*-direction. Indeed, such an auxin gradient controls the *x*-coordinate spot location in such a way that the larger the overall auxin parameter *k*_20_, the more spots are likely to occur. However, as was discussed in section 4, there are instabilities that occur when an extra spatial dimension is present. In other words, when an RH cell is modelled as a two-dimensional flat and oblong cell, certain pattern formation attributes become relevant that are not present in a 1-D setting. In particular, 1-D stripes generically break up into localized spots, which are then subject to possible secondary instabilities. We now explore the effect that a 2-D spatial distribution of auxin has on such localized ROP spots.

### 5.1 A 2-D Auxin Gradient

Next, we perform a numerical bifurcation analysis of the RD system (1.2) for an auxin distribution of the form given in (ii) of (1.1). Such a distribution represents a decreasing concentration of auxin in the *X*-direction, as is biologically expected, but with a greater longitudinal concentration of auxin along the midline *Y* = *L*_*y*_*/*2 of the flat rectangular cell than at the edges *Y* = 0, *L*_*y*_.

To perform a numerical bifurcation study we discretize (1.2) using centered finite differences and we adapt the 2-D continuation code written in MATLAB given in [34]. We compute branches of steady-state solutions of this system using pseudo arclength continuation, and the stability of these solutions is computed a posteriori using MATLAB eigenvalue routines. The resulting bifurcation diagram is shown in Figure 11(a). We observe that there are similarities between this bifurcation diagram and the one for the 1-D problem shown in Figure 6 of [6] in that only fold-point bifurcations were found for the branches depicted. As similar to bifurcation diagrams associated with homoclinic snaking theory on a finite domain (cf. [3, 5, 7, 14, 30]), we observe a key distinguishing feature: solutions belong to a single branch of steady states, undergoing a sequence of fold bifurcations and, in some cases, a change in stability. In large intervals of *k*_20_ we observe multistability which, in turn, indicates that hysteretic transitions between solutions with varying number of spots can occur.

**Figure 11:**
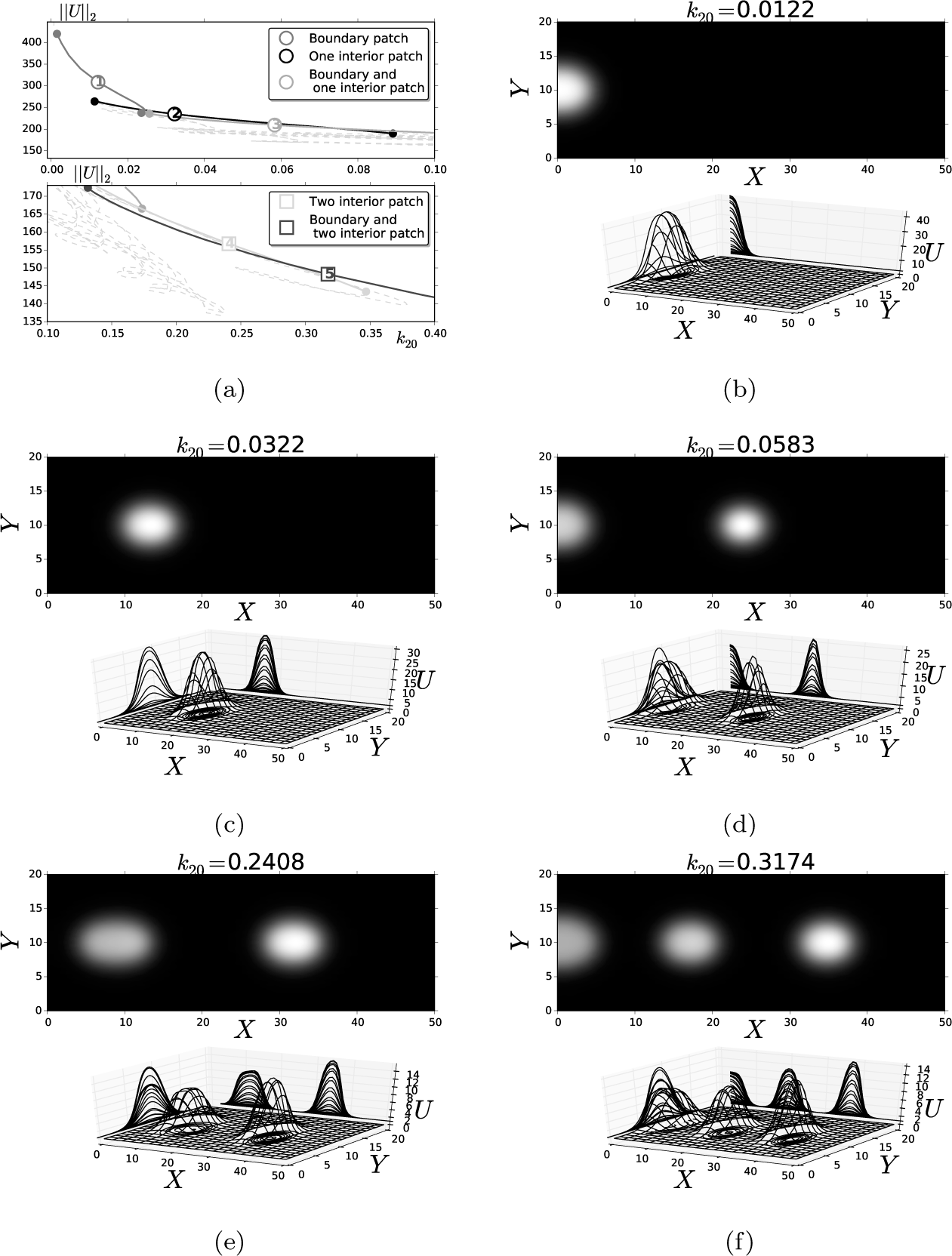
*Bifurcation diagram of the RD system* (1.2) *in terms of the original parameters while varying k*_20_. *Here α* = *α* (**X**) *as given in (ii) of* (1.1) *in a 2-D-rectangular domain. (a) Stable branches are the solid lines, and filled circles represent fold points; top and bottom panels show overlapping of branches of each steady solution: (b) boundary spot, (c) one interior spot, (d) boundary and one interior spot, (e) two interior spots, and (f) boundary and two interior spots. Parameter Set B, as given in Table 1, was used. The k*_20_ *values are given on top of the upper panels for each steady-state in (b)-(f).*

To explore fine details of this bifurcation behavior, the bifurcation diagram in Figure 11(a) has been split into two parts, with the top and bottom panels for lower and higher values of *k*_20_, respectively. From Figure 11(b) we observe that a boundary spot emerges for very low values of the overall auxin rate. This corresponds to branch 1 of Figure 11(a). As *k*_20_ is increased, stability is lost at a fold-point bifurcation which gives rise to branch 2 of Figure 11(a), which gathers a family of single interior spot steady-states, as shown in Figure 11(c). As *k*_20_ increases further, this interior spot solution persists until stability is lost at a fold-point bifurcation. Branch 3 in Figure 11(a) consists of stable steady-states of one interior spot and one boundary spot, as shown in Figure 11(d). Furthermore, Figure 11(e) shows that steady-states consisting of two interior spots occur at even larger values of *k*_20_. This corresponds to branch 4 of Figure 11(a). Finally, at much larger values of *k*_20_, Figure 11(f) shows that an additional spot is formed at the domain boundary. This qualitative behavior associated with increasing *k*_20_ continues, and leads to a creation-annihilation cascade similar to that observed for 1-D pulses and stripes in [6] and [4], respectively. In other words, overlapping stable branches yield steady-states consisting of interior and/or boundary spots that appear or disappear as *k*_20_ is either slowly increased or decreased, respectively.

### 5.2 Instabilities with a 2-D Spatially-Dependent Auxin Gradient

Similar to the 1-D studies in [4, 6], the auxin gradient controls the location and number of 2-D localized regions of active ROP. As the level of auxin increases in the cell, an increasing number of active ROPs are formed, and their spatial locations are controlled by the spatial gradient of the auxin distribution. Moreover, in analogy with the theory of homoclinic snaking, overlapping of stable solution branches occurs, and this leads to a wide range of different observable steady-states in the RD system. These signature features of the bifurcation structure suggest that homoclinic snaking can occur for (1.2), similar to that observed in [5] for a RD system with spatially homogeneous coefficients. In passing, we note that the snaking observed in our system is slanted. Slanted snaking has been reported in systems with conserved quantities (see for instance [10, 36]). The system under consideration does not have a conserved quantity, and the slanting is caused by the auxin gradient. In other words, the gradient gives rise to a strongly slanted homoclinic snaking behavior.

Transitions from unstable to stable branches are determined through fold bifurcations, and are controlled by 𝒪 (1) time-scale instabilities. To illustrate this behavior, we perform a direct numerical simulation of the full RD model (1.2) taking as initial condition an unstable steady-state solution consisting of two rather closely spaced spots, as shown in Figure 12(a). The time-evolution of this initial condition is shown in the different panels of Figure 12. We first observe from Figure 12(b) that these two spots begin to merge at the left domain boundary. As this merging process continues, a new-born stripe emerges near the right-hand boundary, as shown in Figure 12(c). This interior stripe is weakest at the top and bottom boundaries, due to the relatively low levels of auxin in these regions. In Figure 12(d), the interior stripe is observed to give rise to a transversally aligned peanut structure, which remains centered at the *Y*-midline for the same reasons as above at the top and bottom boundaries, whilst another peanut structure in the longitudinal direction occurs near the left-hand boundary. The right-most peanut structure slowly collapses to a solitary spot, while the left-most form undergoes a breakup instability, yielding two distinct spots, as shown in Figure 12(e). Finally, Figure 12(f) shows a steady-state solution with three localized spots that are spatially aligned along the *Y*-midline.

**Figure 12:**
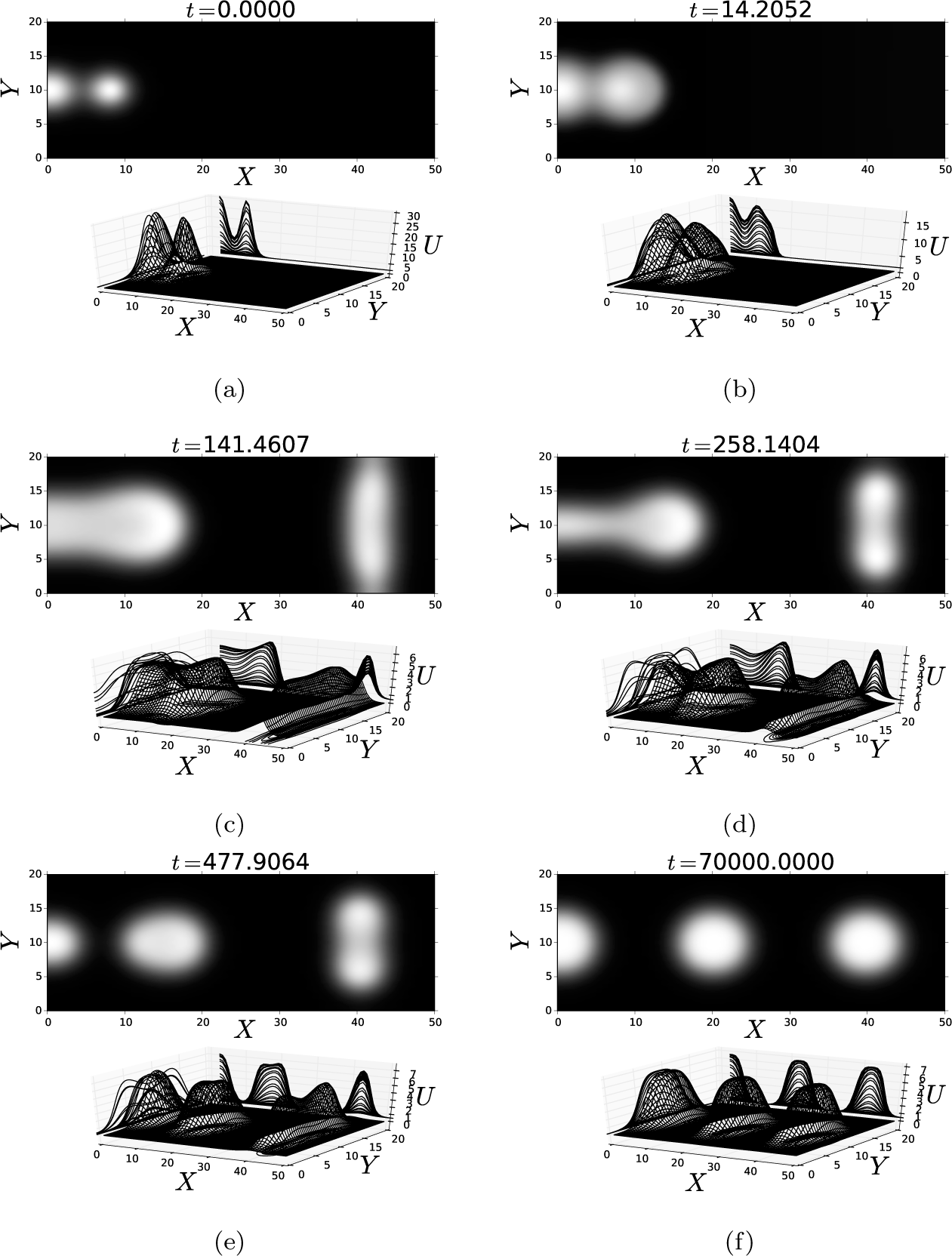
*Numerical simulations of the full RD system with the 2-D spatial auxin distribution given by type (ii) of* (1.1). *The initial condition, shown in (a), is a boundary and interior spot unstable steady-state. The subsequent time-evolution of this steady-state is: (b) spots merging; (c) a new homoclinic stripe is born; (d) a peanut structure emerges; (e) an interior spot arises from a collapsed peanut structure, and (f) finally, a two interior and one boundary spot stable steady-state. Parameter Set B, as given in Table 1 with k*_20_ = 0.0209, *was used.*

In the instabilities and dynamics described above, the 2-D auxin gradient apparently play a very important role, and leads to ROP activation in the cell primarily where the auxin distribution is higher, rather than at the transversal boundaries. In particular, although a homoclinic-type transverse interior stripe is formed, it quickly collapses near the top and bottom domain boundaries where the auxin gradient is weakest. This leads to a peanut structure centered near the *Y*-midline, which then does not undergo a self-replication process but instead leads to the merging, or aggregation, of the peanut structure into a solitary spot. The transverse component of the 2-D auxin distribution is essential to this behavior.

We remark that the asymptotic analysis in section 2 and section 3 for the existence and slow dynamics of 2-D quasi steady-state localized spot solutions can also readily be implemented for the 2-D auxin gradient type (ii) of (1.1). The self-replication threshold in subsection 4.1 also applies to a 2-D spatial gradient for auxin. However, for an auxin gradient that depends on both the longitudinal and transverse directions, it is difficult to analytically construct a 1-D quasi steady-state stripe solution such as that done in [6]. As a result, for a 2-D auxin gradient, it is challenging to obtain a dispersion relation predicting the linear stability properties of a homoclinic stripe (see for instance [12, 24]).

## 6. Discussion

We have used a combination of asymptotic analysis, numerical bifurcation theory, and full time-dependent simulations of the governing RD system to study localized 2-D patterns of active ROP patches, referred to as spots, in a two-component RD model for the initiation of RH in the plant cell *Arabidopsis thaliana*. This 2-D study complements our previous analyses in [4, 6] of corresponding 1-D localized patterns. In our RD model, where the 2-D domain is a projection of a 3-D cell membrane and cell wall, as shown in Figure 2, the amount of available plant hormone auxin has been shown to play a key role in both the formation and number of 2-D localized ROP patches observed, while the spatial gradient of the auxin distribution is shown to lead to the alignment of these patches along the transversal midline of the cell. The interaction between localized active ROP patches and our specific 2-D domain geometry is mediated by the inactive ROP concentration, which has a long range diffusion. This long-range spatial interaction can trigger the formation of localized patches of ROP, which are then bound to the cell membrane. In addition, as is illustrated in Figure 7, our results in Proposition 2 suggest that once ROP patches consisting of localized multi-spots are quickly formed, their location is controlled by the auxin gradient and RH cell shape, where the latter seems to play a more relevant part than in a less realistic 1-D setting (cf. [6]). In this way, a sensitive interplay between overall auxin levels and geometrical properties is crucial to regulate separation of active-ROP patches, which promotes the onset of RHs in mutants. This may qualitatively describe findings on the *rhd6* mutation of *Arabidopsis thaliana* reported in [29].

From a mathematical viewpoint, our hybrid asymptotic-numerical analysis of the existence and slow dynamics of localized active ROP patches, in the absence of any time-scale instabilities, extends the previous analyses of [9, 24, 25, 35] of spot patterns for prototypical RD systems by allowing for a spatially-dependent coefficient in the nonlinear kinetics, representing the spatially heterogeneous auxin distribution. In particular, we have derived an explicit DAE system for the slow dynamics of active ROP patches that includes typical Green’s interaction terms that appear in other models (cf. [9, 24, 25, 35]), but that now includes a new term in the dynamics associated with the spatial gradient of the auxin distribution. This new term contributing to spot dynamics is shown to lead to a spot-pinning effect (cf. [32]) whereby localized active ROP patches become aligned along the transverse midline of the RH cell. We have also determined a specific criterion that predicts when an active ROP patch will become linearly unstable to a shape deformation. This criterion, formulated in terms of the source parameter *S*_*j*_ for an individual spot encodes the domain geometry and parameters associated with the inactive ROP density. In addition, the effect of auxin on instability and shape of active-ROP patches is captured by this parameter, and the nonlinear algebraic system for the collection of source parameters encodes the spatial interaction between patches via a Green’s matrix. When *S*_*j*_ exceeds a threshold, the predicted linear instability of peanut-splitting type is shown to lead to a nonlinear spot self-replication event.

Although our asymptotic analysis of localized active ROP patches has been developed for an arbitrary spatial distribution of auxin, we have focused our study to two specific, biologically reasonable, forms for the auxin distribution given in (1.1). The first form assumes that the auxin concentration is monotone decreasing in the longitudinal direction, as considered in [6] and [4] and motivated experimentally, with no spatially transverse component. The second form is again monotone decreasing in the longitudinal direction, but allows for a spatially transverse component. This second form is motivated by the fact that our 2-D model is a projection of a 3-D cell membrane and cell wall. As a result, it is biologically plausible that some auxin can diffuse out of the transverse boundaries of our 2-D projected domain, leading to lower auxin levels near the transverse boundaries than along the midline of the cell.

We performed a numerical bifurcation analysis on the full RD model for both specific forms for auxin to study how the level of auxin determines both the number of ROP patches that will appear and the increasing complexity of the spatial pattern. This study showed similar features as in the 1-D studies in [4] in that there are stable solution branches that overlap (see Figure 5(a) and Figure 11(a)), which allows for hysteretic transitions between various 2-D pattern types through fold-point bifurcations. In addition, owing to a homoclinic snaking type structure, a creation-annihilation cascade yields 2-D spatial patterns where both spots, baby droplets, and peanut structures can occur. Starting from unstable steady-states, our full time-dependent simulations have shown that 𝒪(1) time-scale competition and self-replication instabilities can occur leading to transitions between various spatial patterns. For the auxin distribution model with a transverse spatial dependence, we showed numerically that the active ROP patches are more confined to the cell midline and that there are no active ROP homoclinic stripes.

As an extension to our analysis, it would be worthwhile to consider a more realistic domain geometry rather than the projected 2-D geometry considered herein. In particular, it would be interesting to study the effect of transversal curvature on the dynamics and stability of localized active ROP patches. In addition, in [4, 6] and here, transport dynamics of ROPs have been modeled under the assumption of a standard Brownian diffusion process. Even though the experimental observations of pinning in [13, 15] seem to reproduced, at least qualitatively, by our model, it would be interesting to extend our model to account for, possibly, more realistic diffusive processes. In particular, it would be interesting to consider a hyperbolic-type diffusive process having a finite propagation speed for signals, as governed by the Maxwell–Cattaneo transfer law (cf. [28, 39]), or to allow for anomalous diffusion (cf. [19]).

## Acknowledgments

D. A., V. F. B.–M. and M. J. W. were supported by EPSRC grant EP/P510993/1 (United Kingdom), UNAM–PAPIIT grant IA104316–RA104316 (Mexico) and NSERC Discovery Grant 81541 (Canada), respectively.

